# N-acetyltransferase 10-mediated mRNA N4-acetylation is Essential for the Translational Regulation During Oocyte Meiotic Maturation in Mice

**DOI:** 10.1101/2024.03.15.585321

**Authors:** Lu Chen, Shao-Yuan Liu, Rui-Bao Su, Yu-Ke Wu, Wen-Jing Wang, Xuan Wu, Song-Ying Zhang, Jie Qiao, Qian-Qian Sha, Heng-Yu Fan

**Affiliations:** MOE Key Laboratory for Biosystems Homeostasis and Protection and Innovation Center for Cell Signaling Network, Life Sciences Institute, Zhejiang University, Hangzhou 310058, China; Assisted Reproduction Unit, Department of Obstetrics and Gynecology, Sir Run Run Shaw Hospital, School of Medicine, Zhejiang University, Hangzhou 310016, China; Fertility Preservation Laboratory, Reproductive Medicine Center, Guangdong Second Provincial General Hospital, 510317 Guangzhou, China; State Key Laboratory of Female Fertility Promotion, Center for Reproductive Medicine, Department of Obstetrics and Gynecology, Peking University Third Hospital, Beijing 100191, China; Center for Biomedical Research, Shaoxing Institute, Zhejiang University, Shaoxing 312000, China

**Keywords:** female fertility, germ cell, RNA modification, transcriptome, translation, maternal-to-zygotic transition

## Abstract

Mammalian oocyte maturation is driven by the strict translational regulation of maternal mRNAs stored in the cytoplasm. However, the function and mechanism of post-transcriptional chemical modifications, especially the newly identified N4-acetylcytidine (ac^4^C) modification catalyzed by N-acetyltransferase 10 (NAT10), are unknown. In this study, we developed a low-input ac^4^C sequencing technology, ac^4^C LACE-seq, and mapped 8241 ac^4^C peaks at the whole-transcriptome level using 50 mouse oocytes at the germinal vesicle stage. We profiled the mRNA landscapes of NAT10-interactions and ac^4^C modifications. The NAT10-interacted and ac^4^C-modified transcripts are associated with high translation efficiency in oocytes. Oocyte-specific *Nat10* knockout wiped out ac^4^C signals in oocytes and caused severe defects in meiotic maturation and female infertility. ac^4^C LACE-seq results indicated that *Nat10* deletion led to a failure of ac^4^C deposition on mRNAs encoding key maternal factors, such as MSY2, ZAR1, BTG4, and cyclin B1, which regulate transcriptome stability and maternal-to-zygotic transition. *Nat10*-deleted oocytes showed decreased mRNA translation efficiency during meiotic maturation, partially due to the direct inhibition of ac^4^C sites on specific transcripts. In summary, we developed a low-input, high-sensitivity mRNA ac^4^C profiling approach and highlighted the important physiological function of ac^4^C in the precise regulation of oocyte meiotic maturation by enhancing translation efficiency.

## Introduction

Mammalian oocyte maturation is driven by the strictly regulated polyadenylation and translational activation of maternal mRNA stored in the cytoplasm. Many mRNAs in germinal vesicle (GV)-arrested oocytes are stored within ribonucleoproteins that protect them from degradation ^1^. Selective polyadenylation and decapping are the major control mechanisms that lead to translational regulation, storage, and degradation ^2,3^. Mouse oocytes are an ideal model for studying the regulation of post-transcriptional cytoplasmic mRNA polyadenylation and translation because fully grown mammalian oocytes are transcriptionally quiescent, and meiosis is driven by a translation product of mRNA stored in the cytoplasm during early development ^4^. Little is known about how mRNA post-transcriptional modifications precisely regulate the translation process during oocyte meiotic maturation, and their specific physiological functions remain largely unknown.

With the innovation and development of high-throughput sequencing technology, "epi-transcriptome" research has gradually emerged in recent years and has become a hot topic in life sciences in the post-genomic era. The “epi-transcriptome” consists of the chemical modifications that occur on the ribonucleotides of RNA after transcription ^5^. More than 170 chemical modifications have been discovered in the RNA of prokaryotes, archaea, and eukaryotes ^6–8^. To date, 11 chemical modifications have been detected in the cytosine nucleoside of RNA, including three types: 5-methylcytidine (m^5^C), 5-hydroxymethylcytidine (hm^5^C), and N4-acetylcytidine (ac^4^C), which are conserved in all living species ^6^. In recent years, an increasing number of studies have found that post-transcriptional modifications of mRNA are involved in various precise regulatory processes, such as RNA transport and output, splicing and processing, polyadenylation, stability, and degradation, to maintain mRNA turnover in organisms ^5^.

N4-acetylcytidine (ac^4^C) was first discovered in yeast tRNA ^9^, and further studies found it in yeast tRNA^Leu^ ^10^, E. coli tRNA^Met^ ^11^, and bacterial tRNA^Met^ ^12^ and revealed that ac^4^C modification of tRNA^Met^ is critical for coding accuracy in protein synthesis^13^. Subsequently, researchers detected ac^4^C in eukaryotic 18sRNA ^14^. In 2018, a study detected ac^4^C modifications in the mRNA of human HeLa cells for the first time and proved that the ac^4^C modification of mRNA was also catalyzed by N-acetyltransferase 10 (NAT10). Further research has shown that ac^4^C modification enhances mRNA stability and promotes translation efficiency ^15^. However, another study concluded that the ac^4^C modification does not occur in human or yeast mRNA. Supporting evidence is that the research group has developed the transcriptome sequencing technology ac^4^C-Seq at a single-nucleotide resolution level. This technology cannot be used to directly detect ac4C modifications in human and yeast mRNA; however, many ac^4^C modifications can be directly detected in archaeal mRNA without induction ^16^. Recently, we reported dynamic changes in the overall ac^4^C modification abundance in total RNA of different tissues and during spermatogenesis. Male germ cell-specific *Nat10* knockout disrupts the normal transcriptome of spermatogenic cells and leads to sterility in male mice, indicating that NAT10-mediated ac^4^C modification has crucial physiological functions ^17^. However, the distribution of ac^4^C modifications in the mRNA during oocyte maturation and the molecular mechanisms regulating this process remain unclear.

To date, researchers have developed various techniques to qualitatively or quantitatively detect ac^4^C modifications in organisms, including high-performance liquid chromatography ^18^, reversed-phase high-performance liquid chromatography ^19^, liquid chromatography-tandem mass spectrometry ^20^, and capillary electrophoresis ^21^. Although these methods can detect ac^4^C modifications to a certain extent, they cannot be used to conduct quantitative studies on the distribution of ac^4^C. "Epi-transcriptome" research in the post-genomic era relies on the innovation and development of various high-throughput sequencing technologies. In recent years, with the development of ac^4^C-specific antibodies ^22^, researchers have developed acRIP-seq based on ac^4^C antibodies^15^. Although the above detection methods have promoted ac^4^C-related research to a certain extent, they cannot be used to study the precise distribution pattern and dynamic change process of ac^4^C modification in the transcriptome, which greatly limits detailed studies of the physiological functions of ac^4^C modification. Over the past two years, researchers have discovered that sodium borohydride and its derivatives can reduce the ac^4^C modification of RNA under acidic conditions. Mismatches occur at sites where reduction reactions occur during reverse transcription. Combined with Sanger sequencing, the specific locations and numbers of mismatches can be detected. This chemical reaction-induced mutation combined with the sequencing technology ac^4^C-Seq can accurately detect the distribution of ac^4^C modifications in the transcriptome at the single-nucleotide resolution level^16,23–25^. However, this method is similar to acRIP-seq in that the library construction process relies on a higher input of samples, which greatly limits the application of these two methods to microvolume samples, such as germ cells and early embryos. Therefore, there is an urgent need to develop ac^4^C sequencing technology with high-sensitivity, single-nucleotide resolution, and low-input amounts.

Recently, researchers have developed LACE-seq to identify RNA-binding protein (RBP) targets in low-input samples, including oocytes. In this approach, the RBP-binding site is directly obtained by linearly amplifying the termination signal of the reverse transcriptase at the RBP-binding site. The elution process was eliminated to reduce the loss of samples, thereby achieving precise identification of RBP-binding sites at single-base resolution and the microcell level ^26^.

In this study, we optimized and improved the key steps of the previously reported LACE-seq method and established a low-input, high-sensitivity ac^4^C LACE-seq technology suitable for oocyte and embryo ac^4^C profiling. We mapped ac^4^C peaks at the whole-transcriptome level in mouse oocytes at the GV stage and constructed an oocyte-specific *Nat10* knockout mouse model to study the physiological function of ac^4^C modification mediated by NAT10 during oocyte maturation. These studies revealed that *Nat10* deletion led to a failure to establish ac^4^C modification of key genes functional in meiotic maturation and further disrupted oocyte maturation-associated mRNA translation activity.

## Results

### NAT10 is expressed in oocytes and associated with the dynamic changes in the abundance of ac^4^C

To investigate the potential function of NAT10 in mediating RNA ac^4^C modification in mouse oocytes during growth and meiotic maturation, we detected the expression of *Nat10* mRNA and protein. RT-qPCR results showed that the expression level of *Nat10* transcripts was high at the GV stage and then continued to decrease until the lowest expression level was observed at the 2-cell stage (Fig. 1A). Western blotting results also indicated that the expression of NAT10 is high in GV oocytes and gradually decreased during the maternal-zygotic transition process (Fig. 1B). Immunofluorescence staining revealed that NAT10 was mainly localized around the nucleolus and nucleoplasm of GV oocytes. Following meiotic resumption and GV breakdown, NAT10 was diffusely distributed in the ooplasm at metaphases I and II (MII). NAT10 gradually accumulated in the pronuclei after fertilization (Fig. 1C). Recent studies have detected RNA chemical modifications, such as m^6^A^27–29^, m^5^C^30^, and ac^4^C^31^ through immunofluorescence staining. Therefore, to detect the presence of RNA ac^4^C modifications in mouse oocytes and preimplantation embryos, we performed immunofluorescence staining using an ac^4^C antibody. The results showed that the ac^4^C signal was present in the nucleus and cytoplasm of oocytes and preimplantation embryos, with the strongest signal in the nucleolus of fully grown GV oocytes (Fig. 1D). Ribosomal RNA is the most abundant RNA in cells and previous studies have confirmed that ac^4^C modifications occur in rRNA. This explains the significant accumulation of the ac^4^C signal in the nucleolus. Following the resumption of meiosis in oocytes, ac^4^C signals are diffused in the ooplasm. After fertilization, ac^4^C signals relocalized to nucleolus-like structures (Fig. 1D), which was consistent with the expression pattern of the NAT10 protein. These results indicate that NAT10 is dynamically expressed in mouse oocytes. Its expression pattern was consistent with the changes in ac^4^C localization and levels, suggesting that NAT10 plays a crucial role in regulating RNA ac^4^C modification during oogenesis.

**Figure 1.**
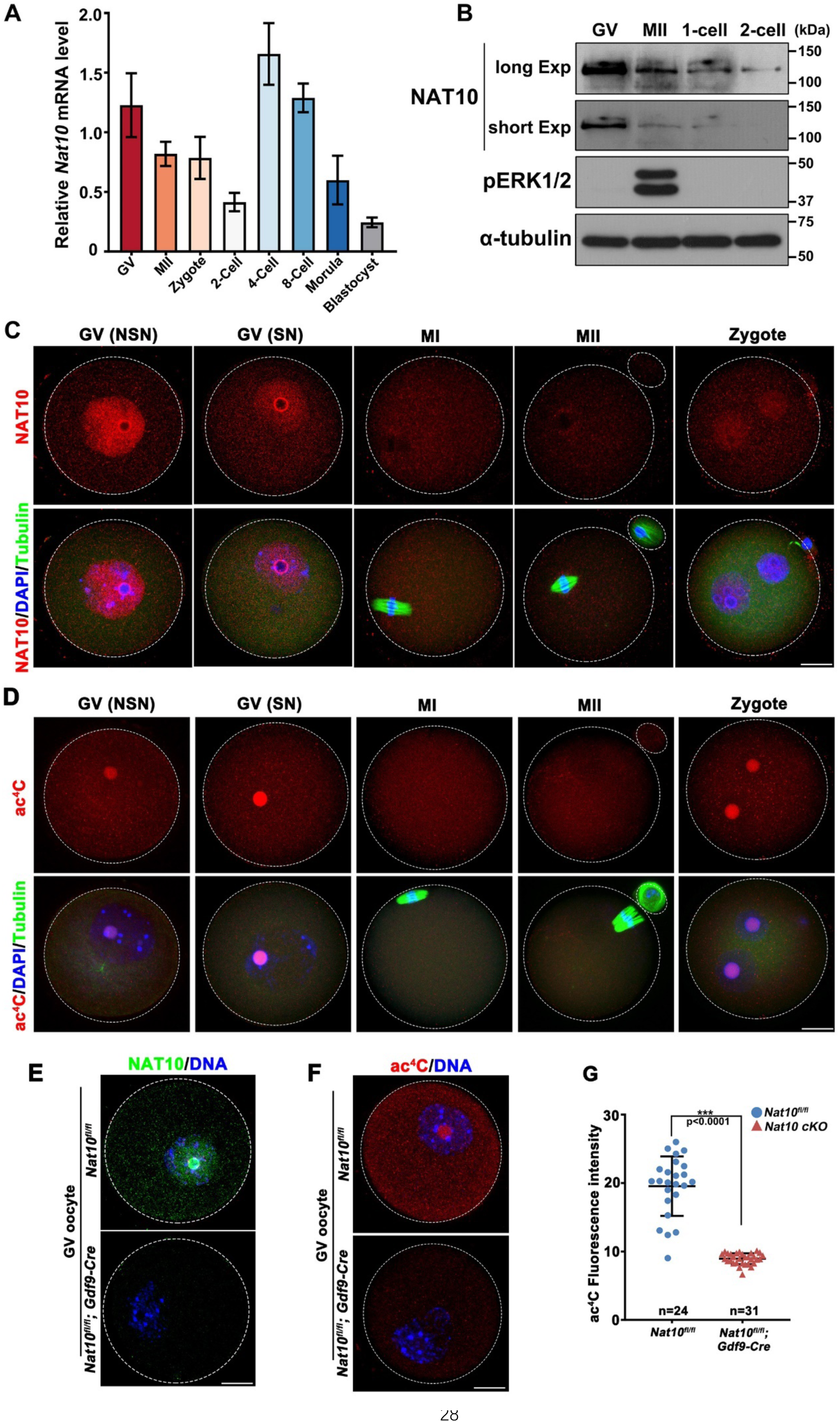
Expression and localization of NAT10 in oocyte and preimplantation embryos. **A:** RT-qPCR results showing the level changes of *Nat10* transcripts during mouse oocyte maturation and preimplantation embryonic development. **B:** The expression of NAT10 during the maternal-to-zygotic transition stages was detected using western blotting. **C–D:** Immunofluorescence results showing the expression and localization of NAT10 and ac^4^C during mouse oocyte meiotic maturation and fertilization. **E–F:** Immunofluorescence results showing the expression and localization of NAT10 and ac^4^C in WT and *Nat10*-deleted oocytes. **G:** Quantitative statistics of ac^4^C fluorescence signal in (E). Scale bars = 20 μm for all panels in **C**, **D**, and **E**.

To confirm whether the deletion of *Nat10* in oocytes caused a reduction in the overall RNA ac^4^C modification level, we collected GV stage oocytes from wild-type (WT) and *Nat10^fl/fl^;Gdf9-Cre* mice, which are characterized in detail in Figure 4, and performed NAT10 and ac^4^C immunofluorescence staining. The results showed that the NAT10 signal was undetectable in oocytes from *Nat10^fl/fl^;Gdf9-Cre* mice (Fig. 1E) and that the ac^4^C signals in *Nat10-*deleted oocytes were significantly reduced (Fig. 1F–G), proving that the RNA ac^4^C modifications detected in WT oocytes were mediated by NAT10. This result also confirmed that the NAT10 and ac^4^C signals detected using immunofluorescence were specific.

### LACE-seq results show ac^4^C modifications on transcripts in mouse oocytes

In this study, LACE-seq was improved. The ac^4^C antibody was directly cross-linked with RNA to produce a steric hindrance effect, and the same linear amplification reverse transcriptase formed a termination signal, thereby obtaining a specific modification of ac^4^C at the whole-transcriptome level. The technical route is shown in Fig. 2A and the Materials and Methods section. To further confirm the credibility of the ac^4^C peaks captured by LACE-seq, we performed LACE-seq on transcripts interacting with NAT10, the only known ac^4^C writer protein.

**Figure 2.**
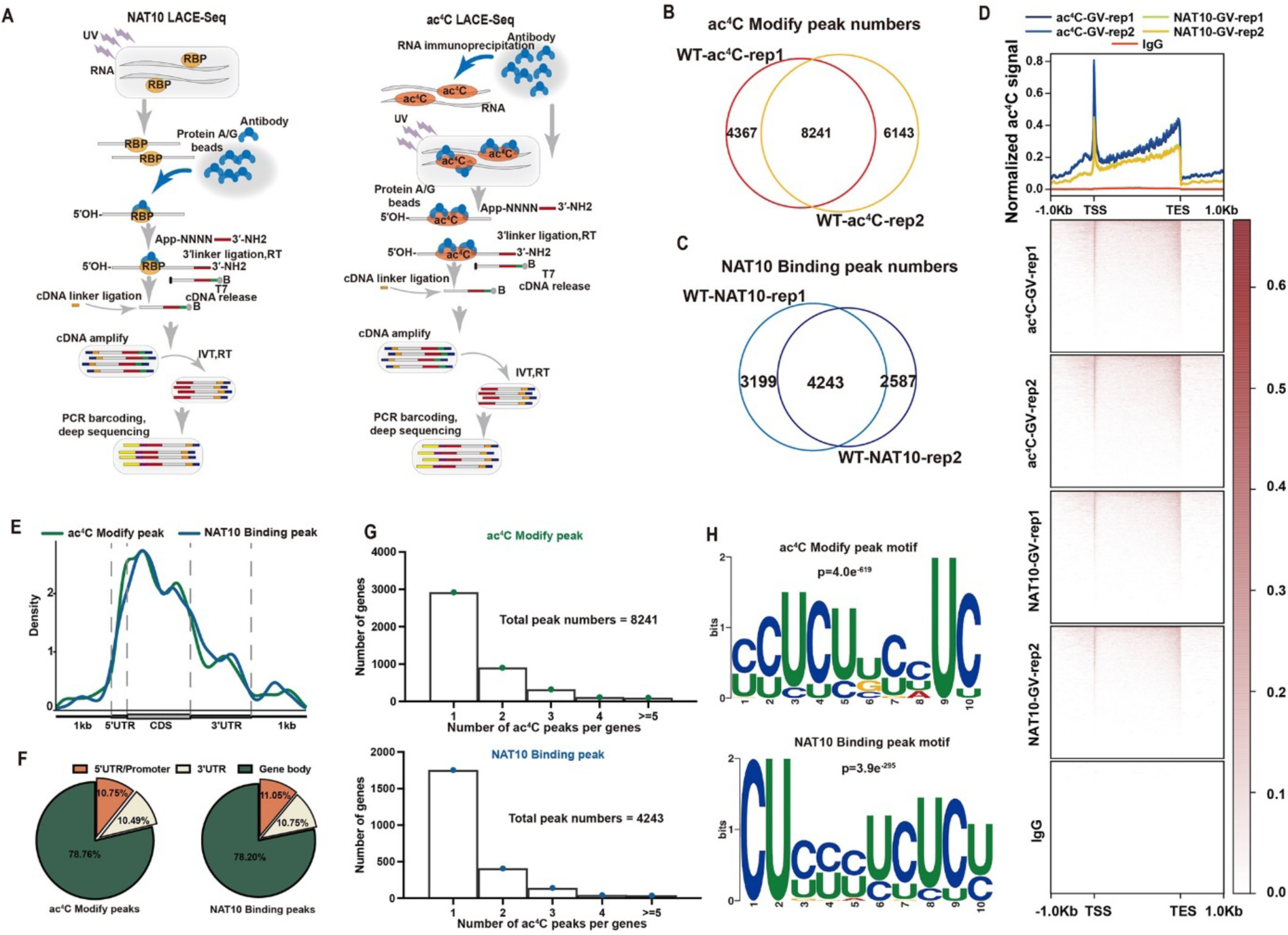
Identification of ac^4^C on maternal transcripts in mouse oocytes using a LACE-seq-based approach. **A:** Technical diagram of NAT10 LACE-seq (left) and ac^4^C LACE-seq (right). **B:** Overlap of ac^4^C peaks from two independent experiments in WT GV oocytes. **C:** Overlap of NAT10-binding peaks from two independent experiments in WT GV oocytes. **D:** Distributions and heatmap of ac^4^C and NAT10-binding peaks on the genome. IgG served as a control. TSS: transcription start site; TES: transcription end site. **E:** ac^4^C-modified and NAT10-RNA interacting sites were enriched around the mRNA. **F:** Distribution ratio of ac^4^C and NAT10-binding peaks in each transcript region. **G:** Histogram showing the average number of NAT10 and ac^4^C-binding peaks per gene in WT GV oocyte. **H:** Motif enrichment analyses of ac^4^C and NAT10-binding peaks.

Based on this optimized ac^4^C LACE-seq method, we collected 50 WT GV oocytes for ac^4^C and NAT10 LACE-seq library construction, and two replicates were set for each group of samples. The results showed that the quality of the data obtained at this starting amount was highly reliable, and the correlation between the two biological replicates in the WT GV oocyte NAT10 LACE-seq and ac^4^C LACE-seq results was 80% (Fig. S1A and Supplementary Table S1). The overlap ratio between the two biological repeats was high (Fig. 2B-C). We considered the overlapping peaks of biological repeats as credible ac^4^C peaks (8241) and NAT10-binding peaks (4243) (Fig. 2B–C and Supplementary Table S2-S7). In comparison, no ac^4^C or NAT10 signals were detected in the IgG group, indicating that the peaks captured by the ac^4^C and NAT10 antibodies were specific (Fig. 3F). Further analyses indicated that the ac^4^C peaks and NAT10 binding peaks had consistent distribution patterns in the genome, with higher signals enriched near the transcription start signal (TSS) and transcription ending signal (TES) (Fig. 2D). The peaks detected by both ac^4^C and NAT10 antibodies mainly distributed in the coding sequenced and 5′ untranslated regions (UTR) (Fig. 2E–F). Further analysis of the 8241 ac^4^C and 4243 NAT10-binding peaks showed that most detected transcripts carry 1–2 peaks (Fig. 2G). The motifs of ac^4^C-and NAT10-binding peaks were CU-enriched (Fig. 2H), which is consistent with the previously reported distribution pattern and motif characteristics of ac^4^C ^15,16^. These results further verify that the ac^4^C signal obtained by ac^4^C LACE-seq has high sensitivity and reliability.

**Figure 3.**
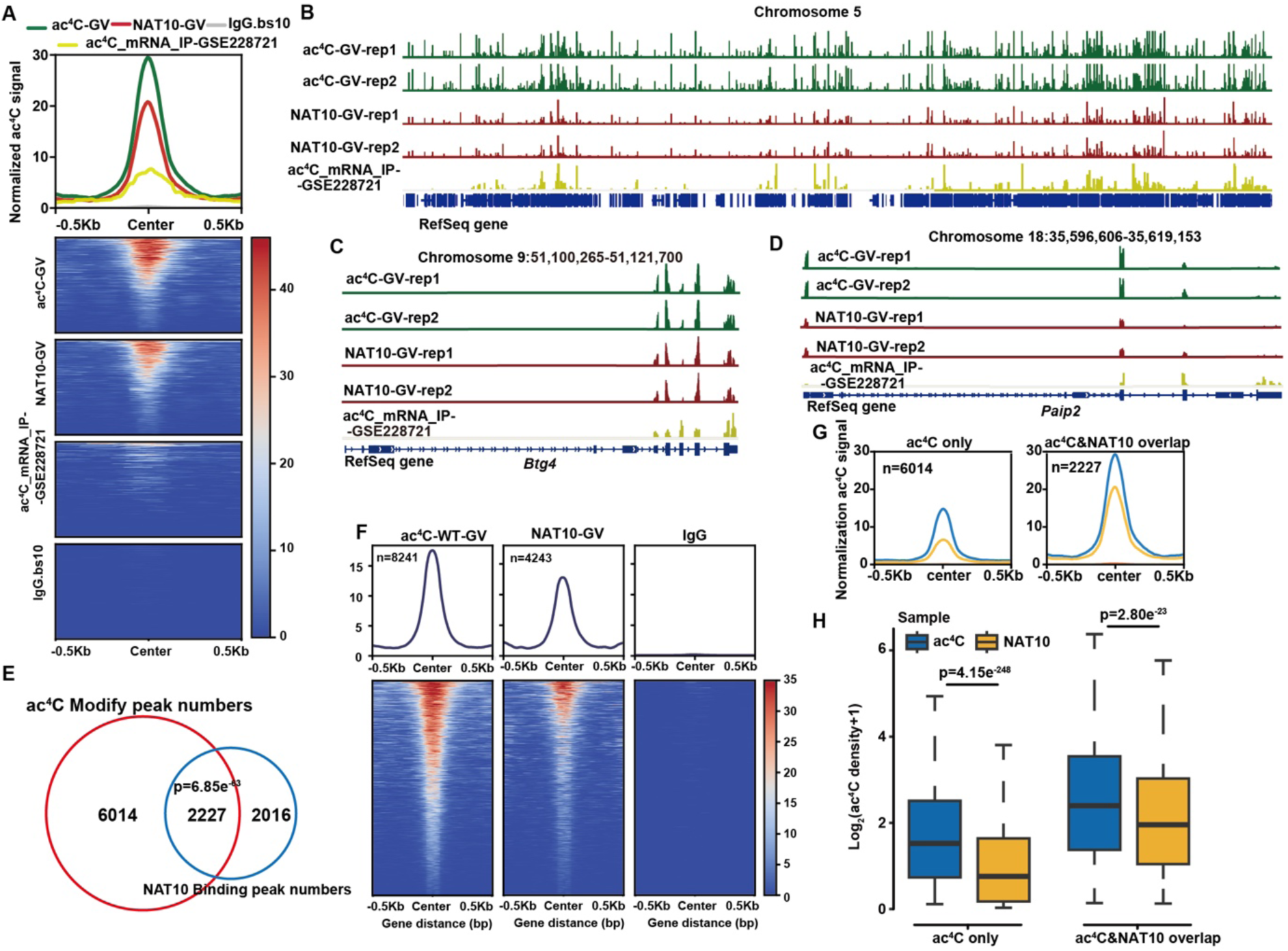
Comparative analyses of oocyte ac^4^C LACE-seq and oocyte acRIP-seq. **A:** Metaprofile and heatmap of signals from ac^4^C LACE-seq, NAT10 LACE-seq, and acRIP-seq (GSE228721) around the identified peaks. IgG served as a control. **B:** UCSC genome browser views of ac^4^C LACE-seq, NAT10 LACE-seq, and acRIP-seq reads on chromosome 5. **C–D:** UCSC genome browser views of distributions of peaks obtained by ac^4^C LACE-seq, NAT10 LACE-seq, and acRIP-seq of *Btg4* (C), and *Paid2* (D), both are key oocyte genes. **E:** The overlapping targeted peaks by ac^4^C LACE-seq and NAT10 LACE-seq in GV stage oocytes. **F:** The heat map shows the peak signals captured by ac^4^C LACE-seq and NAT10 LACE-seq in GV oocytes, and the IgG group is set as a negative control. **G:** The ac^4^C peaks signal intensity normalization analysis was performed on the peaks captured only by ac^4^C LACE-seq and the peaks signals co-targeted by ac^4^C and NAT10. **H:** Boxplot showing that the NAT10 targets have a higher ratio of ac^4^C occupancy than NAT10 to binding sites. The P-value was calculated by one-tailed unpaired Student’s *t*-test. The center line represents the median, the box borders represent the first and third quartiles, and the whiskers are the most extreme data points within 1.5 × the interquartile range.

**Figure 4.**
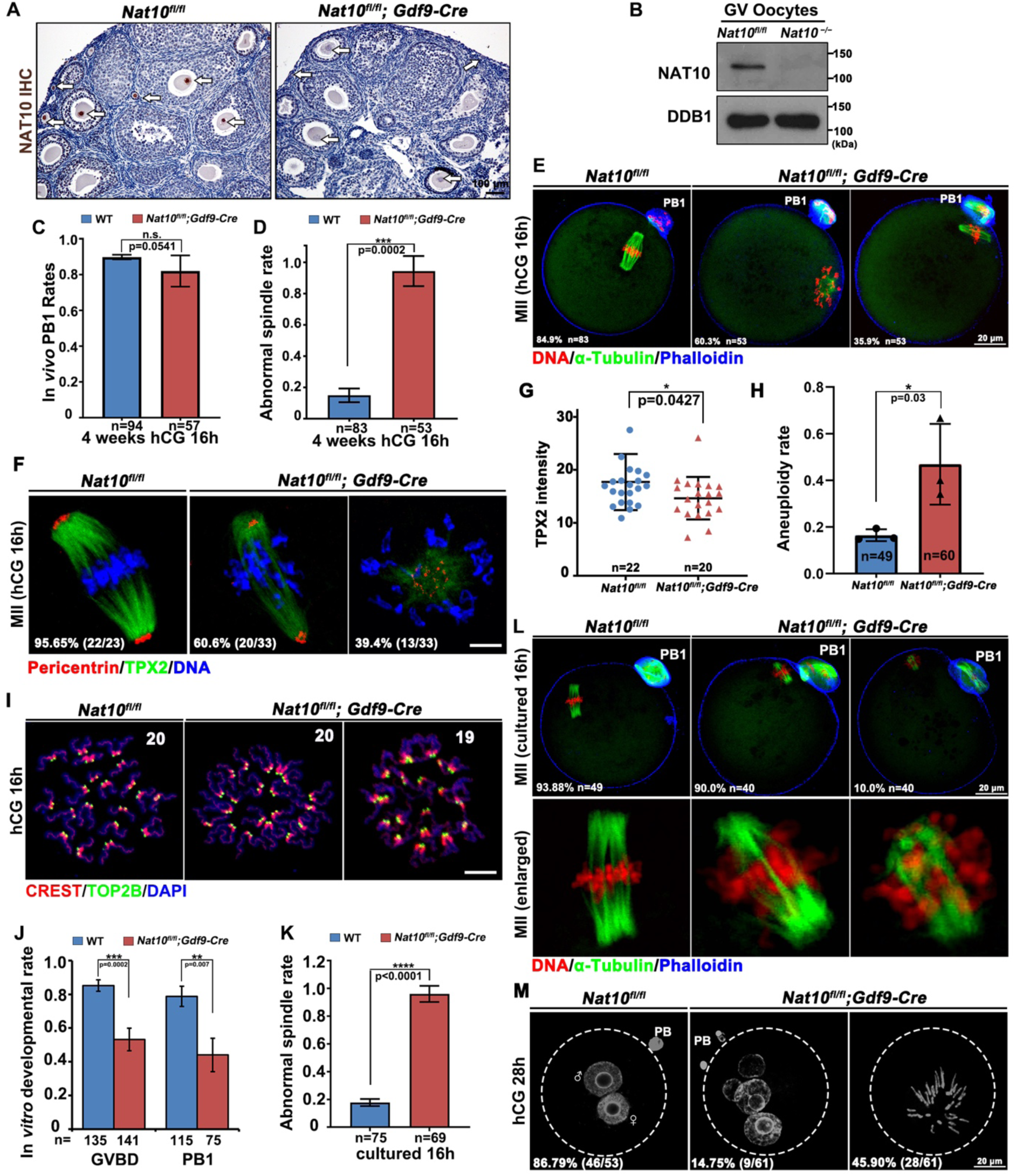
*Nat10* deletion results in oocyte maturation defects. **A:** Immunohistochemistry results indicating deletion of NAT10 protein in oocytes within ovarian follicles. Scale bar = 100 μm. **B:** Western blot results showing NAT10 protein levels in oocytes of WT and *Nat10^fl/fl^;Gdf9-Cre* mice. The total proteins of 100 oocytes were loaded in each lane. A constitutively expressed protein DDB1 was blotted as a loading control. **C:** Statistics on the proportion of the first polar body (PB1) emissions in oocytes ovulated by WT and *Nat10^fl/fl^;Gdf9-Cre* mice at 16 h after hCG injection. **D:** Ratio of abnormal spindles in ovulated oocytes. **E:** Immunofluorescence staining to detect spindle assembly and chromosome arrangement in MII oocytes of WT and *Nat10^fl/fl^;Gdf9-Cre* mice. **F:** Immunofluorescence was used to detect the expression and localization of TPX2 and pericentrin proteins in MII stage oocytes ovulated by WT and *Nat10^fl/fl^;Gdf9-Cre* mice. Scale bars = 10 μm. Quantifications of TPX2 immunofluorescence signal intensity in (I). **H:** Statistics on the proportion of chromosomal aneuploidy in MII stage oocytes ovulated by WT and *Nat10^fl/fl^;Gdf9-Cre* mice. **I:** Chromosome spreading combined with immunofluorescence detection of chromosomal euploidy in MII oocytes ovulated by WT and *Nat10^fl/fl^;Gdf9-Cre* mice. Scale bars = 10 μm. **J:** Meiotic resumption (characterized by germinal vesicle breakdown (GVBD)) and PB1 emission rates of WT and *Nat10*-deleted oocytes cultured *in vitro*. **K:** Ratio of abnormal spindles in WT and *Nat10*-deleted oocytes cultured *in vitro*. **L:** Immunofluorescence staining to detect spindle assembly and chromosome arrangement in WT and *Nat10*-deleted oocytes at 16 h after *in vitro* culture. Scale bars = 20 μm. **M:** Detection of pronucleus formation by DAPI immunofluorescence staining in fertilized eggs derived from WT and *Nat10^fl/fl^;Gdf9-Cre* female mice. Scale bars = 20 μm.

### ac^4^C LACE-seq results have good reproducibility with previously published data and in cross analyses

To verify the reliability and reproducibility of the ac^4^C LACE-seq method, we performed joint analyses of ac^4^C LACE-seq data and acRIP-seq results in GV oocytes ^32^. In this study, the authors performed acRIP-seq on 2000 GV stage oocytes and identified 6188 transcripts carrying ac^4^C peaks. In our study, only 50 GV stage oocytes were used, and 8241 ac^4^C peaks (7461 transcripts) were collectively captured in both sets of replicates (Fig. 2B). In addition, ac^4^C signals detected by ac^4^C LACE-seq were stronger than those detected by acRIP-seq (Fig. 3A). Chromosomes 5 and 10 were selected to display the overall ac^4^C peaks using the Integrative Genomics Viewer, and the signals captured by ac^4^C LACE-seq and NAT10 LACE-seq were stronger than those of acRIP-seq (Fig. 3B and Fig. S1B). Signals of ac^4^C modification were detected in transcripts of key maternal genes, including polyadenylate-binding protein 1 (*Pabpn1)*, BTG anti-proliferation factor 4 (*Btg4)*, and poly(A) binding protein interacting protein 2 (*Paip2*), and the results showed that ac^4^C LACE-seq had a stronger sensitivity than acRIP-seq in oocytes at the GV stage (Fig. 3C–D and Fig. S1C). However, the initial input samples for LACE-seq were only 2.5% (50 GV for ac^4^C LACE-seq and 2000 GV for acRIP-seq) of the acRIP-seq. Furthermore, acRIP-seq in oocytes cannot pinpoint the accurate positions of cytosine acetylation owing to the technical limitation of ∼200 bp mRNA fragment resolution, LACE-seq could map ac4C modified sites in 50 oocytes at single-nucleotide resolution. The motifs of ac^4^C-and NAT10-binding peaks were CU-enriched (Fig. 2H), which is consistent with the previously reported distribution pattern and motif characteristics of ac^4^C ^15,16^.

We further compared the results of ac^4^C and NAT10 LACE-seq. Nearly half (2227/4243) of the targets captured by the NAT10 antibody were also captured by ac^4^C LACE-seq (Fig. 3E). Furthermore, the peaks captured by the ac^4^C antibody had more significant signal enrichment in the peak center regions than those captured by the NAT10 antibody (Fig. 3F). This is consistent with our prediction because the ac^4^C antibody directly recognizes the modification site, whereas the NAT10 antibody only targets NAT10-interacting transcript regions mediated by other RNA-binding proteins. The signal intensities of the ac^4^C peaks detected by ac^4^C LACE-seq and NAT10 LACE-seq were significantly higher than the ac^4^C signals detected only by one of the two approaches. Similar to the trend in Fig. 3F, the signal intensities of the peaks captured only by ac^4^C LACE-seq were higher than those captured by NAT10 LACE-seq alone (Fig. 3G). Further quantitative analysis of the ac^4^C signal intensity in these two groups (ac^4^C-only and ac^4^C and NAT10 overlap) showed that the overlapping group had the highest signal intensity of ac^4^C peaks, whereas the NAT10-only group had the lowest signal intensity. In addition, the signal intensity captured by the ac^4^C antibody was significantly higher than that captured by the NAT10 antibody in all three groups (Fig. 2H). Collectively, these results confirm the reliability of the ac^4^C LACE-seq method and indicate that ac^4^C LACE-seq is a low-input, high-sensitivity, and high-reproducibility approach for profiling ac^4^C modifications at the transcriptome level.

### NAT10 is essential for female fertility and oocyte meiotic maturation

*Nat10* knockout mice are embryonic lethal ^33^. To study the *in vivo* function of NAT10, we generated *Nat10^fl/fl^;Gdf9-Cre* conditional knockout mice that specifically inactivate *Nat10* expression in oocytes at the primordial follicle stage. Similar to the immunofluorescence results shown in Fig. 1E, immunohistochemical staining of ovarian paraffin sections of *Nat10^fl/fl^;Gdf9-Cre* mice showed that NAT10 protein was not detected in oocytes at all follicle stages (Fig. 4A). Western blot results indicated that NAT10 was expressed in GV oocytes and downregulated in MII oocytes after meiotic maturation but was undetectable in oocytes from *Nat10^fl/fl^;Gdf9-Cre* mice (Fig. 4B and Fig. 7E). These results demonstrate that *Nat10* is efficiently and specifically inactivated in *Nat10^fl/fl^;Gdf9-Cre* mice. For fertility tests, we crossed *Nat10^fl/fl^;Gdf9-Cre* and *Nat10^fl/fl^* female mice with WT male mice for six months. *Nat10^fl/fl^;Gdf9-Cre* female mice were completely sterile, suggesting that *Nat10* is essential for reproduction in female mice.

To determine the causes of female infertility, we evaluated the *in vivo* maturation of oocytes in WT and *Nat10^fl/fl^;Gdf9-Cre* mice following superovulation treatment. In *Nat10*-deleted oocytes collected from the oviducts, the polar body 1 emission rate was consistent with that in the control group (Fig. 4C). However, immunofluorescence staining of α-tubulin in the ovulated MII oocytes revealed that the proportion of spindle abnormalities significantly increased after *Nat10* deletion (Fig. 4D–E). Immunofluorescence staining of TPX2 and pericentrin, important proteins for spindle assembly, showed that the loss of *Nat10* resulted in disordered spindle assembly and abnormal localization of pericentrin (Fig. 4F), and the fluorescence intensity of TPX2 on the spindles was significantly reduced (Fig. 4G). We collected ovulated MII oocytes and performed chromosomal spreading and immunofluorescence staining. The results showed that *Nat10* deletion increased aneuploidy in MII oocytes (Fig. 4H-I). We further evaluated oocyte meiotic maturation using *in vitro* culture experiments. The results showed that GV breakdown and polar body 1 emission rates were significantly reduced in *Nat10-*deleted oocytes compared with WT oocytes (Fig. 4J). Immunofluorescence staining showed the proportion of spindle abnormalities in MII stage oocytes of *Nat10^fl/fl^;Gdf9-Cre* mice exceeded 90% under *in vitro* culture conditions (Fig. 4K– L).

To evaluate the developmental potential of these oocytes, we performed superovulation on *Nat10^fl/fl^;Gdf9-Cre* female mice, mated them with WT male mice, and collected fertilized eggs from the oviducts. The pronucleus formation rate was significantly reduced after *Nat10* deletion, and the proportion of abnormal pronuclei increased (Fig. 4M). These results further confirm the meiotic maturation defects and reduced fertilization potential of *Nat10*-deficient oocytes.

### Loss of *Nat10* results in reduced global mRNA ac^4^C modification

To study the ac^4^C modification changes in transcripts caused by *Nat10* deletion at the whole-transcriptome level, we collected 50 GV stage oocytes from WT and *Nat10^fl/fl^;Gdf9-Cre* mice for ac^4^C LACE-seq and set up IgG LACE-seq as the control. The results showed that the ac^4^C signal was enriched in the WT oocytes and was higher near the TSS and TES sites (Fig. 5A). When *Nat10* was deleted, the ac^4^C signal in GV stage oocytes was significantly reduced to levels comparable to those in the negative control group (Fig. 5A). The ac^4^C signals were enriched in WT GV oocytes within 0.5 kb upstream and downstream of the putative ac^4^C modification center, with a total of 8241 modification sites. When *Nat10* is deleted, the ac^4^C signal within 0.5 kb upstream and downstream of the ac^4^C modification site is globally reduced. Only 215 ac^4^C modification sites with weak signals were detected (Fig. 5B). The consensus-binding motif among these 215 sites was devoid of cytosines, suggesting that the detected ac^4^C signals were nonspecific background signals (Fig. 5C). Gene ontology analysis results indicated that most of the mRNAs carrying NAT10-dependent ac^4^C modifications encoded proteins involved in post-transcriptional RNA regulation, including mRNA processing, RNA-protein interactions, splicing, translation, and degradation (Fig. 5D). We selected five key maternal mRNAs that accumulated abundantly in oocytes to visualize the ac^4^C peaks through Integrative Genomics Viewer (Fig. 5D). Among these, *Msy2, Pabpn1*, zygote arrest 1 (*Zar1)*, and *Btg4* are involved in regulating mRNA translation and stability. Multiple ac^4^C peaks were detected in these transcripts using both ac^4^C and NAT10 LACE-seq in WT oocytes. However, after *Nat10* knockout, the ac^4^C peaks of these transcripts almost completely disappeared (Fig. 5D). These results further suggest that the ac^4^C peaks detected by ac^4^C LACE-seq in the WT oocytes are accurate and specific because the oocytes from *Nat10*-deleted mice serve as a stringent negative control.

**Figure 5.**
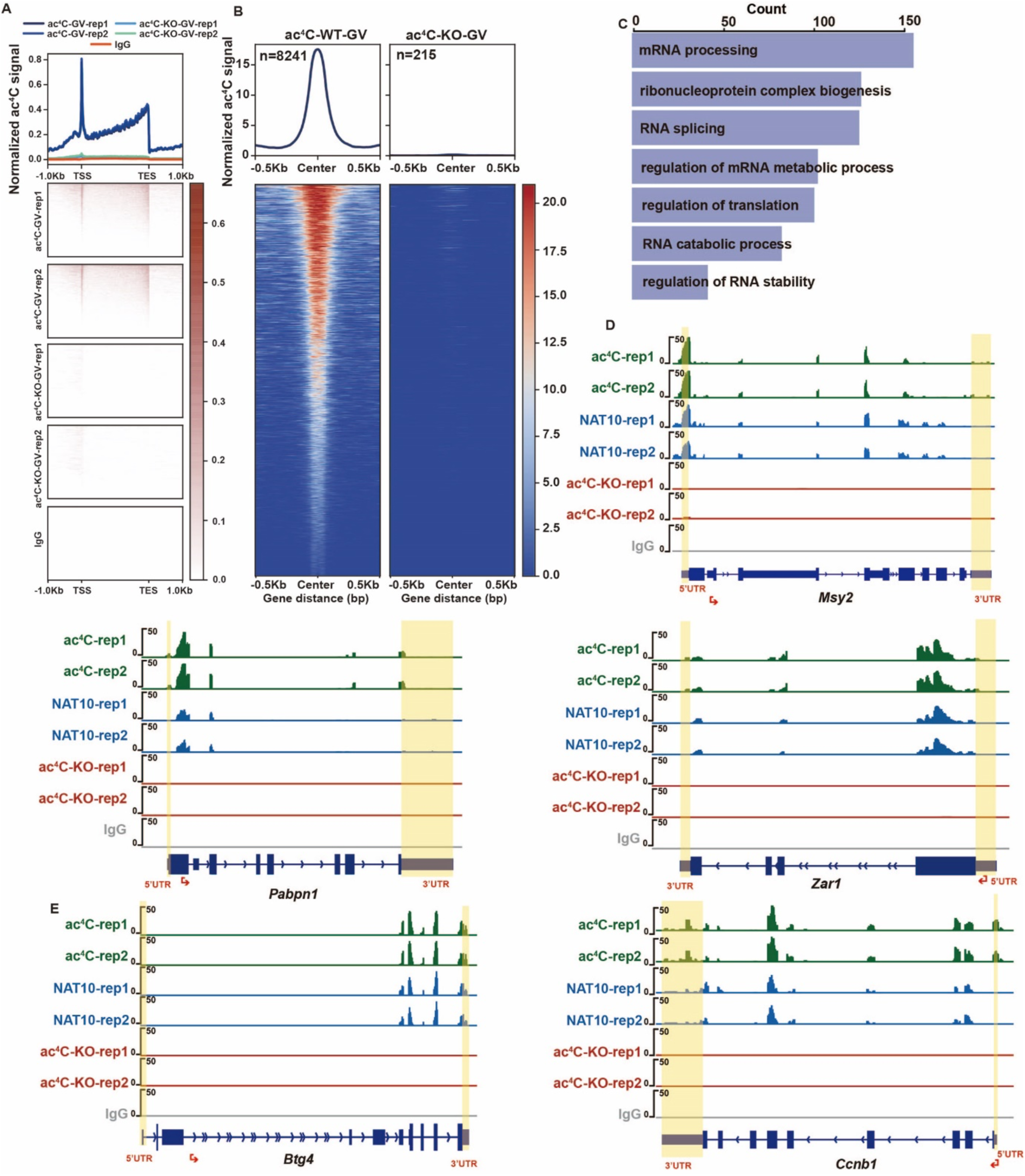
Loss of *Nat10* results in reduced ac^4^C modifications on mRNAs. **A:** ac^4^C LACE-seq detects ac^4^C modification in GV stage oocytes of WT and *Nat10 ^fl/fl^;Gdf9-Cre* mice. **B:** Metaprofile and heatmap of ac^4^C signals around the identified peaks from WT and *Nat10^fl/fl^;Gdf9-Cre* mice. **C:** GO enrichment analysis was performed on genes whose ac^4^C peaks were significantly reduced after *Nat10* deletion. **D:** UCSC genome browser views of abolished ac^4^C LACE-seq reads on the key maternal genes at the GV stage of WT and *Nat10^fl/fl^;Gdf9-Cre* mice.

### Transcripts deregulation caused by *Nat10* deletion is not significantly related to ac^4^C density

To investigate how reduced ac^4^C modification caused by *Nat10* deletion leads to meiotic defects, we collected GV oocytes from WT and *Nat10^fl/fl^;Gdf9-Cre* mice for transcriptome sequencing. RNA-seq results showed high reproducibility among the three biological samples (Fig. S2A). Principal Component Analysis (PCA) was performed on these samples, and the results showed that the principal components between WT and *Nat10*-deleted oocytes were not significantly different (PCA2 = 2.41%) (Fig. S2B), indicating that *Nat10* deletion did not have a significant impact on the overall transcriptome stability. The RNA-seq results were analyzed for differentially expressed genes [Log2(FC) >1 or <-1, P-value <0.05]. The volcano plot shows that the loss of *Nat10* resulted in the downregulation and upregulation of 930 and 1093 transcripts in GV oocytes (Fig. 6A). We further analyzed the relationship between mRNA ac^4^C modifications and the oocyte transcriptome. We defined transcripts carrying ac^4^C peaks with fragments per kilobase million > 1, as well as putative ac^4^C motifs, as ac^4^C^+^ transcripts and used the 2366 defined ac^4^C^+^ transcripts for subsequent analyses (Fig. 6B). Next, we performed an overlap analysis of the differentially expressed genes caused by *Nat10* deletion and the ac^4^C^+^ genes. The results showed that only 59 differentially expressed transcripts were ac^4^C^+^ (Fig. 6C), indicating that NAT10-mediated ac^4^C modification did not significantly affect the stability of ac^4^C^+^ transcripts.

**Figure 6.**
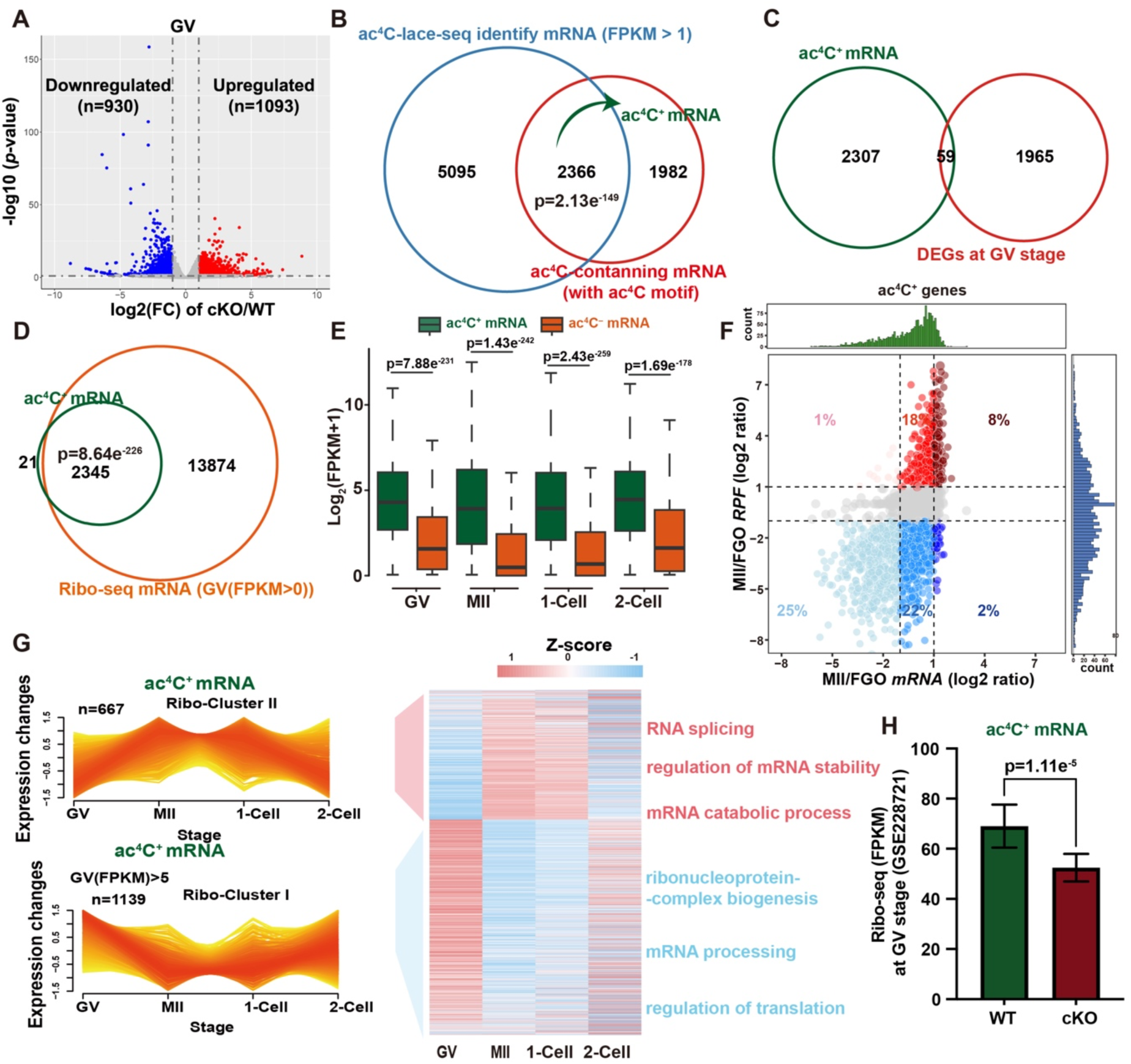
ac^4^C modification abundance couples to translation during oocyte meiotic maturation. **A:** Volcano plots show the number of significantly differentially expressed genes (DEGs) in GV oocytes of WT and *Nat10^fl/fl^;Gdf9-Cre* mice. *P* threshold (= 0.05) and log_2_FC threshold (= ±1) are reported in gray horizontal and vehicle dashed lines, respectively. n: gene number, FC: fold change. **B:** The expression level corresponding to the 7461 transcripts carrying ac^4^C peaks is limited to FPKM > 1, and the transcripts containing ac^4^C motives are defined as ac^4^C^+^ transcripts. **C:** Overlap analysis was conducted between ac^4^C^+^ transcripts (B) and DEGs (A) in GV oocytes with or without *Nat10* knockout. **D:** A Venn diagram showing the overlap between strict ac^4^C^+^ transcripts and ribosome-bound transcripts in GV oocytes. P-values obtained using Fisher’s exact test are indicated. **E:** Boxplot showing Ribo-lite dynamic change of ac4C+ and ac^4^C^-^ transcripts during GV to 2-cell stage. The P-value was calculated using a one-tailed unpaired Student’s *t*-test. The center line represents the median, the box borders represent the first and third quartiles, and the whiskers are the most extreme data points within 1.5× the interquartile range. **F:** Scatter plots showing the ac^4^C^+^ transcripts dynamic changes (fold change, x-axis) and Ribo-lite dynamic change (fold change, y-axis) from GV to MII. Red and blue plots show that translatome upregulated or downregulated more than 2-fold change in the MII stage compared with GV. **G:** 2366 ac^4^C^+^ transcripts were jointly analyzed with Ribo-seq data during the MZT process. They were divided into two clusters based on Ribo-seq expression changes. Heat map showing dynamic changes during MZT. The enriched GO terms are also listed. Histogram showing translatome of the ac^4^C^+^ transcripts in this study by the published Ribo-seq data^32^ in WT and *Nat10*-deleted GV oocytes. The P-value was calculated using one-tailed unpaired Student’s *t*-test. Error bars, S.E.M. **H:** The ac^4^C^+^ gene in this study was jointly analyzed with the published Ribo-seq data^32^ in wild-type and *Nat10* conditional knockout mouse GV oocytes.

**Figure 7.**
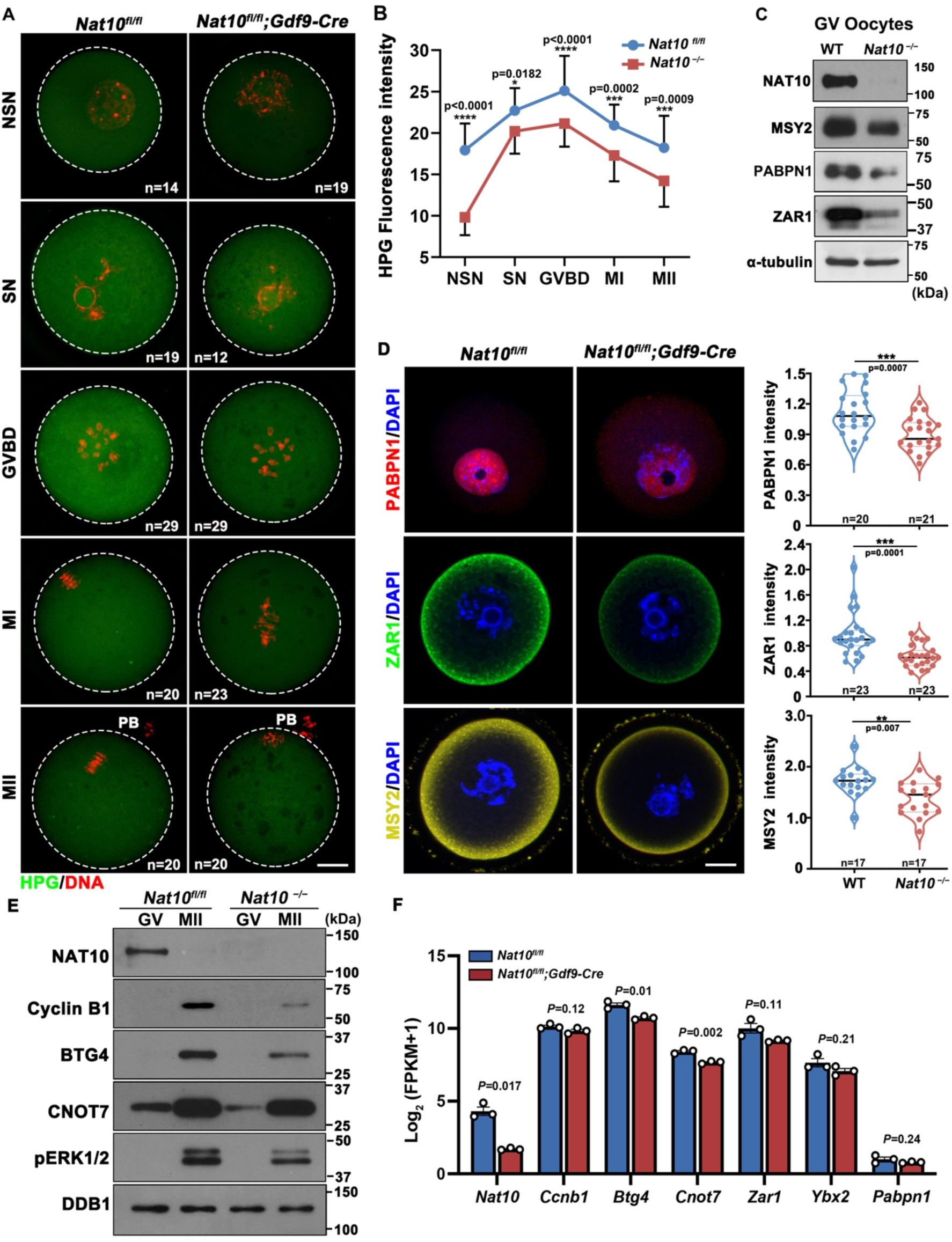
*Nat10* deletion affects mRNA translation efficiency during oocyte maturation. **A:** The Click-iT L-Homopropargylglycine (HPG) Alexa Fluor-488 Protein Synthesis assay detects changes in overall translation levels during oocyte meiotic maturation. Scale bar: 20 μm. **B:** Quantitative statistics of HPG fluorescence signals in (A). **C:** Western blot results showing the levels of the indicated proteins in WT and *Nat10*-deleted GV oocytes. Total proteins of 100 oocytes were loaded in each lane. **D:** Immunofluorescence and quantification results of the indicated proteins in WT and *Nat10*-deleted GV oocytes. Scale bar: 20 μm. **E:** Western blot results showing levels of the indicated proteins in WT and *Nat10*-deleted oocytes before and after meiotic maturation. Total proteins of 100 oocytes were loaded in each lane. **F:** RNA-seq results showing mRNA levels of indicated genes in WT and *Nat10*-deleted oocytes at the GV stage. Scale bars = 20 μm for all panels in **A** and **D**.

Further joint analysis of the ac^4^C^+^ transcripts with the M-decay, Z-decay, and zygotic genome activation-dependent transcripts defined in WT mice showed that 54% and 34% of ac^4^C^+^ transcripts were M-decay and Z-decay transcripts, respectively. Another 12% of transcripts were zygotic genome activation-dependent (Fig. S2C). To analyze the relationship between ac^4^C modification and gene expression, we divided the ac^4^C^+^ transcripts into two clusters based on the changes in expression during the maternal-to-zygotic transition (MZT) in WT mice (Fig. S2D–E). Among them, there were 1113 transcripts in Cluster II, which were gradually degraded during meiotic maturation and then maintained at low levels (Fig. S2E). This suggests that most ac^4^C^+^ transcripts are highly expressed in GV oocytes and may play an important role in meiotic maturation.

### RNA ac^4^C modification couples to translation efficiency during oocyte meiotic maturation

Precise translation regulation is a core biological event that occurs during the meiotic maturation of oocytes. To investigate whether ac^4^C modification is required for oocyte meiotic maturation by affecting translation activity, we compared 2366 ac^4^C^+^ transcripts with translatomes in GV oocytes identified using Ribo-lite ^34^. Almost all ac^4^C^+^ transcripts showed high translational activity in GV oocytes (Fig. 6D). Compared with ac^4^C^-^ transcripts, ac^4^C^+^ transcripts had higher translation activity (ribosome-binding efficiency) at the GV, MII, 1-cell, and 2-cell stages (Fig. 6E). In addition, we analyzed the translational activity of ac^4^C^+^ transcripts in oocytes at the GV and MII stages. The results showed that most ac^4^C^+^ transcripts had higher translational activity in GV than in MII oocytes. However, the mRNA metabolic regulation showed no obvious biases. (Fig. 6F). We divided the ac^4^C^+^ transcripts into two clusters based on changes in their translational activity during the MZT process. In addition, 1139 of them belong to Cluster I, with higher translational activity in the GV stage, and these transcripts are enriched in the biological processes of the ribonucleoprotein complex, mRNA processing, and regulation of translation (Fig. 6G). In comparison, fewer (667) transcripts belonged to Cluster II, with relatively low translational activity at the GV stage (Fig. 6G). These results indicate a positive correlation between mRNA ac^4^C modification and translational activity during oocyte meiotic maturation. To determine the relationship between ac^4^C modification and translation activity further, we analyzed ribosome sequencing results in GV oocytes of WT and *Nat10^fl/fl^;Gdf9-Cre* mice ^32^. The results showed that when *Nat10* was deleted, the translation efficiency (fragments per kilobase million of the Ribo-seq results) of the ac^4^C^+^ transcripts was significantly reduced (Fig. 6G). The inhibition of ac^4^C modification caused by *Nat10* deletion significantly reduces the translation efficiency of NAT10-targeted transcripts, highlighting the coupling of ac^4^C modification and the translation activity of mRNAs.

### *Nat10* deletion and reduced ac^4^C modification impair the translation of key maternal mRNAs

To further verify the impact of *Nat10* deletion on translation activity during the oocyte meiotic maturation, we performed the Click-iT L-Homopropargylglycine (HPG) Protein Synthesis assay on oocytes at multiple stages of meiotic maturation to evaluate the overall translation activity in oocytes. The results showed that translation was rapidly activated as meiosis resumed in WT oocytes. In contrast, translation activity at all detected stages was reduced in *Nat10-*deleted oocytes (Fig. 7A–B). Transcripts encoding key maternal proteins, including MSY2, PABPN1, and ZAR1, underwent ac^4^C modifications at the GV stage, mainly in their coding sequences (Fig. 5F). Both western blotting and immunofluorescence results indicated that the protein expression levels of MSY2, PABPN1, and ZAR1 were compromised in *Nat10*-deleted GV oocytes (Fig. 7C–D). As shown in Fig. 7A, meiotic resumption-coupled translational activation was compromised in *Nat10*-deleted oocytes, we selected key proteins that need to be translationally activated during oocyte meiotic maturation for verification. Western blotting results showed that the expression of BTG4 and cyclin B1 proteins was significantly reduced in *Nat10*-deficient MII oocytes (Fig. 7E). In contrast, the mRNA levels of these key maternal genes were not significantly affected by *Nat10* deletion (Fig. 7F).

To further verify the impact of *Nat10* deletion on meiotic maturation-triggered translational activation, we selected two key maternal genes, *Btg4* and *Ccnb1,* to construct translation reporter plasmids, transcribed the mRNA *in vitro*, microinjected it together with *mCherry* mRNA into GV oocytes from WT and *Nat10^fl/fl^;Gdf9-Cre* mice, and measured translational activity using the ratio of green fluorescent protein (GFP) and mCherry fluorescence intensity. After the injection of GFP-*Btg4*-3’UTR into WT GV oocytes, GFP was rapidly translationally activated after meiosis resumed. Both fluorescence and western blotting results showed that the mRNA translation efficiency of Flag-GFP-*Btg4*-3’UTR was significantly reduced after *Nat10* deletion (Fig. 8A–C). Similarly, the translation activity of Flag-GFP-*Ccnb1*-3’UTR mRNA was significantly reduced in both GV and MII oocytes of *Nat10^fl/fl^;Gdf9-Cre* mice (Fig. 8D–F). These results indicate that *Nat10* deletion affects the translation efficiency of maternal mRNAs in mouse oocytes during meiotic maturation.

**Figure 8.**
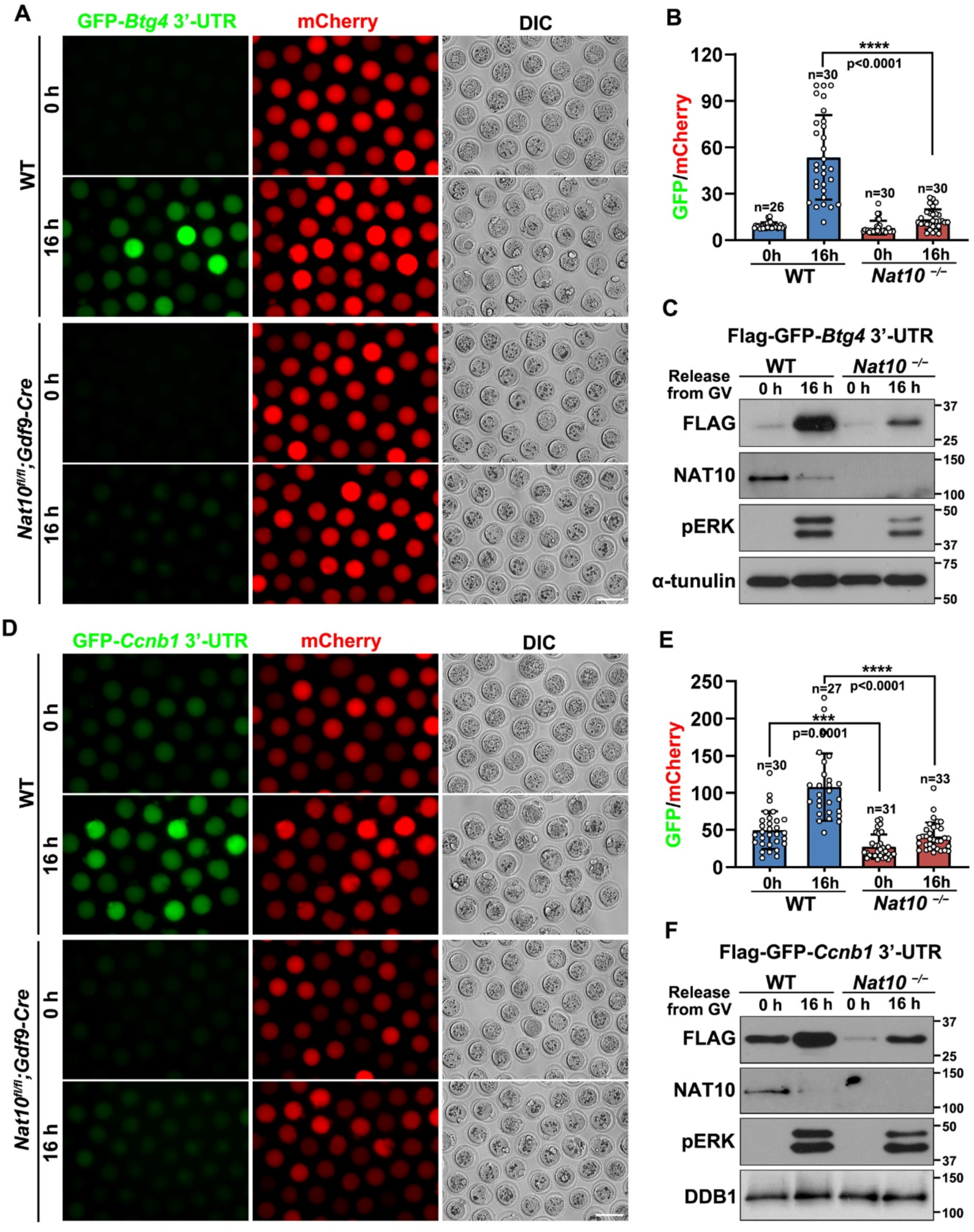
Reduced translational activation of key maternal mRNAs in *Nat10*-deleted oocytes. **A:** *GFP-Btg4 3′-UTR* reporter assay detects *Btg4* mRNA translation efficiency during oocyte maturation in WT and *Nat10^fl/fl^;Gdf9-Cre* mice. **B:** Quantification of GFP/mCherry fluorescence intensity. **C:** Western blotting was used to detect the translational activity of *Btg4 3′-UTR* during oocyte maturation of WT and *Nat10^fl/fl^;Gdf9-Cre* mice. **D:** *GFP-Ccnb1 3′-UTR* reporter assay detects *Ccnb1* mRNA translation efficiency during oocyte maturation in WT and *Nat10^fl/fl^;Gdf9-Cre* mice. **E:** Quantification of GFP/mCherry fluorescence intensity. **F:** Western blotting was used to detect the translational activity of *Ccbn1 3′-UTR* during oocyte maturation of WT and *Nat10^fl/fl^;Gdf9-Cre* mice. Scale bars = 100 μm for all panels in **A** and **D**.

Subsequently, we investigated the direct effects of mRNA ac^4^C modifications on translation. We mutated the ac^4^C modification sites in the coding sequences of *Msy2* and *Zar1* without affecting the amino acid sequences (Fig. 9A). We then transfected 293T cells (which did not endogenously express MSY2 and ZAR1) with plasmids expressing WT and ac^4^C site-mutated *Msy2* and *Zar1*. Western blotting results showed that the ac^4^C site mutation reduced the expression of MSY2 and ZAR1 (Fig. 9B–D). Oocyte microinjections of ac^4^C site-mutated *Msy2* and *Zar1* mRNAs were not performed because NAT10 is localized to the nucleus and cannot catalyze ac^4^C modifications in mRNAs microinjected into the ooplasm.

**Figure 9.**
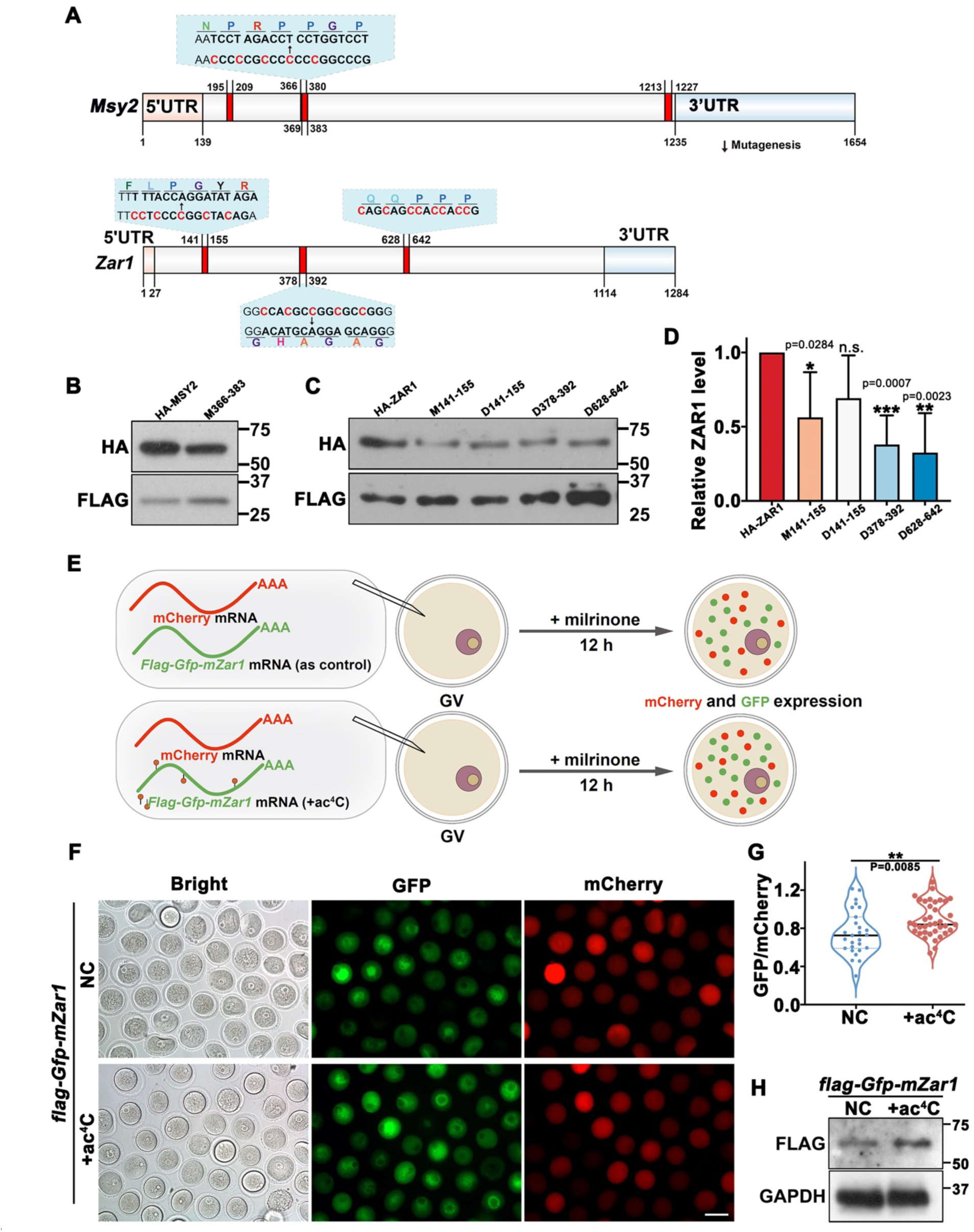
Effects of mRNA ac^4^C modifications on the translation of key proteins in oocytes. **A:** Localization of ac^4^C modification sites in the encoding regions of *Msy2* and *Zar1* transcripts. Selected ac^4^C sites were mutated without affecting the amino acids being encoded. **B–C**: Western blotting results showing protein expression levels of MSY2 and ZAR1 before and after mutations of ac^4^C sites. 293T cells were transfected with plasmids encoding WT and ac^4^C site-mutated *Msy2* and *Zar1* cDNAs. **D:** Quantifications of the band intensities in (C). **E:** Illustration of *in vitro* transcription and oocyte microinjection of *Zar1* mRNAs with or without ac^4^C modifications. **F:** Epifluorescence results show the expression of *Zar1* mRNAs with or without ac^4^C modifications in GV oocytes at 24 h after microinjections. mRNAs encoding mCherry was coinjected as an internal control. Scale bars = 100 μm. **G:** Quantifications of relative GFP/mCherry intensities in (F). **H:** Western blotting results showing expression levels of FLAG-GFP-ZAR1 proteins from *Zar1* mRNAs with or without ac^4^C modifications.

We also microinjected GV stage-arrested oocytes with *in vitro* transcribed GFP-ZAR1 mRNAs with or without ac^4^C incorporation (Fig. 9E). The results showed that transcripts containing ac^4^C had higher translational levels than those without the ac^4^C modification (Fig. 9F–H). These results provide evidence that the ac^4^C modification of mRNAs enhances translation in oocytes.

## Discussion

In this field, the controversy remains about whether ac^4^C modification occurs on eukaryotic mRNA. Arango et al. used antibody-based methods (acRIP-Seq) to map ac^4^C and showed the presence of 4251 acetylated regions in HeLa cells ^15^. In contrast, Sas-Chen et al. developed a quantitative and single-nucleotide resolution approach (ac^4^C-Seq) for the quantitative mapping of ac^4^C sites transcriptome-wide, and the profiling results suggested that ac^4^C is absent or present at a very low stoichiometry in human cell lines and yeast mRNA ^16^. The discrepancy between these two studies is likely due to the differences in detection methodologies. Another reason for this contradiction is that ac^4^C has different modification abundances in different cell lines and tissues. Methods based on antibody enrichment combined with next-generation sequencing have been applied to detect various RNA modifications, such as m^6^A, m^5^C, m^1^A, Ψ, and ac^4^C. The accuracy of the detection results of this method depends on antibody specificity. It is necessary to carefully exclude false-positive results caused by antibody cross-reactivity; therefore, the reproducibility and specificity of these chemically modified peaks obtained by sequencing must be fully verified. However, the limitation of the antibody-based method is that it is impossible to identify where the modification occurred at single-nucleotide resolution. In contrast, ac^4^C-Seq is a single-nucleotide resolution approach that relies on chemical treatment to change the physicochemical properties of modified bases, resulting in mismatches during the subsequent reverse transcription of library construction. However, this approach is limited to detecting low stoichiometric bases in mRNAs expressed at low levels. This method also requires a deeper sequencing depth; otherwise, it is difficult to fully reflect the true stoichiometric sites and the abundance of modifications. In this study, we developed an ac^4^C LACE-seq technology based on ac^4^C antibodies and used this technology to identify 8241 ac^4^C peaks in oocytes. These peaks were abolished after *Nat10* deletion, proving the reliability of this method. Our study provides new evidence for ac^4^C modification in mammalian mRNAs and establishes a LACE-seq technology-based ac^4^C detection method, which will facilitate the study of ac^4^C modification in a broad field.

To verify the ac^4^C LACE-seq method, we also performed LACE-seq of NAT10. NAT10 is the only ac^4^C writer protein currently known. Although NAT10 contains a tRNA-binding domain^35^, it has not been reported to directly bind to mRNAs. The ac^4^C peaks captured by NAT10 are presumably to indirectly bind with NAT10 through its "Adaptor" proteins. To date, RNA-binding proteins that mediate ac^4^C modifications in mRNAs remain elusive. In our study, the ac^4^C peaks captured by both ac^4^C LACE-seq and NAT10 LACE-seq showed higher signal intensities, indicating the credibility of the ac^4^C peaks. Many ac^4^C peaks were captured by ac^4^C LACE-seq than by NAT10 LACE-seq. A likely explanation is that the transcripts dissociated from NAT10 after ac^4^C establishment.

Previously, ac^4^C-related research was limited to the application of low-input detection technologies. In this study, we developed an ac^4^C modification detection technology-ac^4^C LACE-seq based on LACE-seq, which is suitable for a small number of samples. Using this technology, we identified 8241 ac^4^C modification sites on 7461 transcripts in 50 GV stage oocytes. This technology has the advantages of high sensitivity and low input compared with other published ac^4^C detection technologies. The development of this technology will help promote ac^4^C-related research in germ cells and early embryonic development, as well as benefit other fields. In addition, the development of ac^4^C LACE-seq technology will provide new inspiration and a paradigm for developing other mRNA modification detection technologies, arousing widespread interest in epi-transcriptomics.

Using oocyte-specific *Nat10* knockout mice as a model, this study confirmed that ac^4^C modification of mRNA in oocytes is catalyzed by NAT10 and that NAT10-mediated ac^4^C modification has important physiological functions during meiotic maturation and female fertility (Fig. 10). Transcriptome analyses indicated that ac^4^C modification in oocytes is highly coupled to translation efficiency rather than affecting transcript stability and that ac^4^C drives precise regulation of the oocyte meiotic maturation process by enhancing translation efficiency (Fig. 9). In addition to mRNAs, ac^4^C modifications were detected in the tRNAs of leucine and serine and at two sites on the 18S rRNA. We also observed high levels of ac^4^C signals in the nuclear-like bodies of GV oocytes and zygotes, where rRNA was enriched. When *Nat10* was deleted, these nucleolar ac^4^C signals disappeared, confirming that NAT10 was responsible for rRNA ac^4^C modifications in mouse oocytes. It is conceivable that NAT10-mediated ac^4^C modifications of mRNAs, tRNAs, and 18S rRNAs contribute to efficient mRNA translation activity in oocytes. Studies in yeast and cultured mammalian cell lines have shown that the disruption of ac^4^C modifications on tRNAs (tRNA-Ser and tRNA-Leu) and 18S rRNAs delayed the cell growth rate but did not cause cell death, suggesting that ac^4^C modifications on tRNAs and 18S rRNAs facilitate mRNA translation, but are not essential^13,15,36–39^. This observation was consistent with the limited distribution of ac^4^C modifications in tRNAs and 18S rRNAs. In contrast, we provide experimental evidence that ac^4^C modifications of key maternal mRNAs positively regulate their translational efficiencies. We propose that NAT10-mediated ac^4^C modifications in all three types of RNAs, including mRNAs encoding key maternal factors, are crucial for efficient maternal mRNA translation during mouse oocyte maturation (Fig. 10).

**Figure 10.**
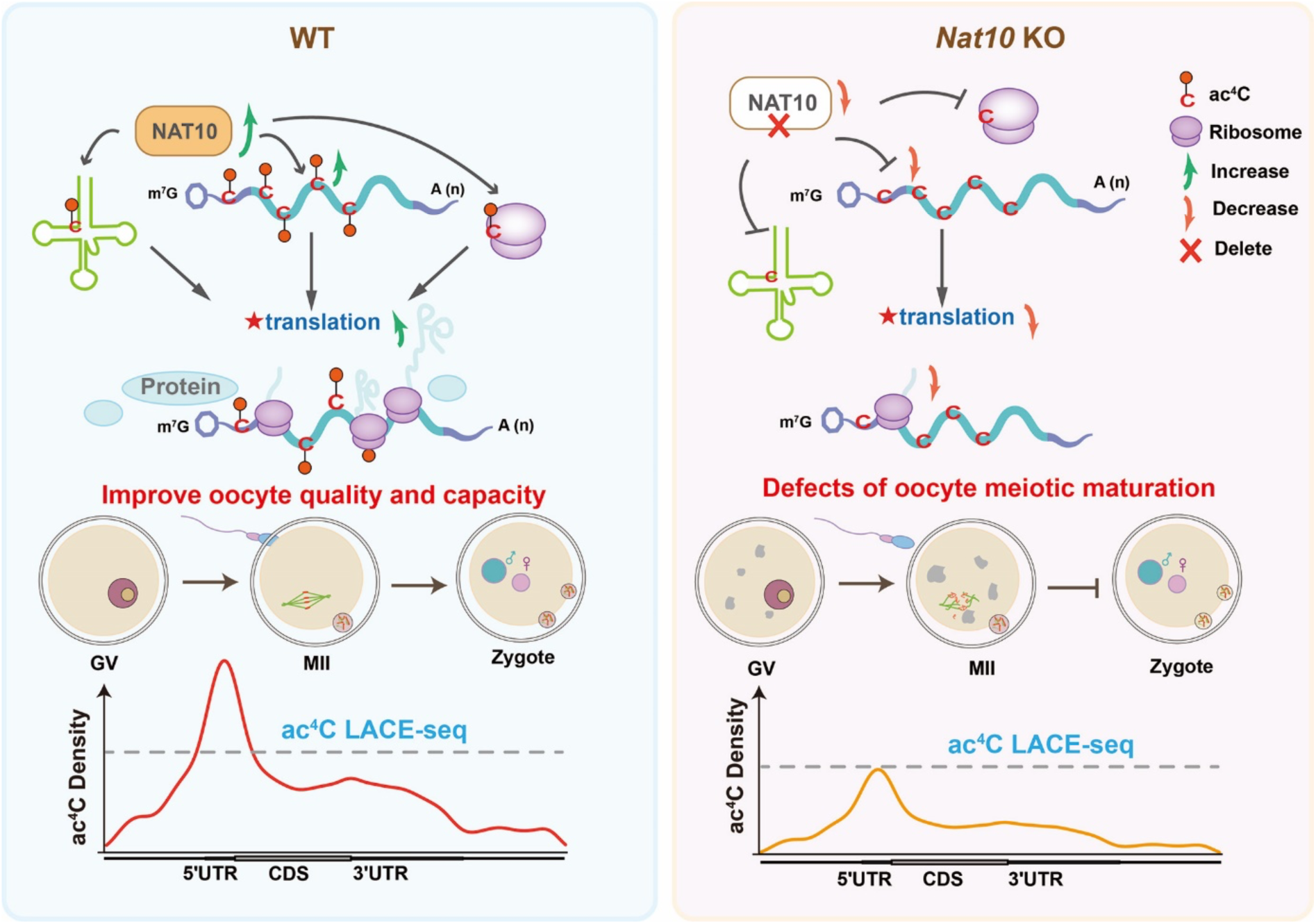
Schematic diagram of NAT10-mediated mRNA N4-acetylcytidine regulates translation of key proteins during oocyte meiotic maturation. In WT mice, NAT10 catalyzes ac^4^C modification on mRNA, tRNAs, and rRNAs and promotes the timely translation of proteins from these key transcripts to ensure normal oocyte meiotic maturation. When NAT10 is absent, the RNA ac^4^C modification cannot be established, and the translation efficiency of these transcripts is reduced, leading to defects in the meiotic maturation process, ultimately leading to infertility in female mice.

In addition to causing mRNA translation defects, *Nat10* deletion also influenced the levels of certain transcripts in oocytes, as shown in our study and a recent report^32^. Ac^4^C modifications may directly affect the stability of some mRNAs; however, the underlying mechanisms remain unknown. In our opinion, *Nat10* deletion in oocytes is likely to affect transcript stability via the following indirect mechanisms: 1) compromised translation of mRNAs encoding key maternal factors involved in transcript storage, particularly MSY2 and ZAR1. Previous studies have shown that *Msy2* or *Zar1* knockout leads to premature decay of maternal mRNAs and defects in oocyte maturation. 2) Impaired translational activation of transcripts encoding proteins responsible for mRNA degradation, such as BTG4 and CNOT7 (this study and Bao et al ^32^.). 3) Ribosome binding and active translation of transcripts also play a role in mRNA stability in mouse oocytes and zygotes. Therefore, ac^4^C modification may indirectly affect the stability of maternal mRNAs by regulating their translation efficiency.

We acknowledge the limitations of this study. We have not yet mapped the dynamic changes in ac^4^C modification during the entire meiotic maturation process and preimplantation embryonic development and failed to provide Ribo-seq data at each stage owing to technical limitations. We attempted to identify ac^4^C-modified mRNAs in MII oocytes and zygotes using the ac^4^C LACE-seq method without success. Maternal mRNAs undergo extensive degradation during meiotic maturation and fertilization^40,41^. However, de novo transcription did not occur during this process. Therefore, it is possible that the levels of ac^4^C-modified maternal mRNAs at these developmental stages were too low to be effectively detected using the ac^4^C LACE-seq method.

Currently, the mechanisms through which mRNA ac^4^C modifications facilitate translation are unknown. Studies in cultured somatic cells have suggested that ac^4^C modifications increase the binding efficiency of transcripts to ribosomes, especially at the wobbled third base of codons^13,15,42^. Interactions between mRNA ac^4^C modification sites and key RNA-binding proteins, particularly those involved in the translational regulation of oocytes, remain to be investigated.

## Materials and Methods

### Animals

All experiments involving mouse strains used in this study involved a C57BL/6J genetic background. The *Gdf9-Cre* transgenic mouse line has been previously described ^43^. *Nat10*-floxed mice (*Nat10 ^flox/flox^*) in which exons 2 and 3 were flanked with sites, as previously reported^17^, were crossed with the *Gdf9-Cre* mouse line to generate *Nat10* conditional knockout mice. Mice were housed in cages under the specific pathogen-free environment of the Laboratory Animal Center of Zhejiang University, which controlled at 50–70% humidity, a 12/12-h light/dark cycle with 20–22 °C temperature, and water and food provided ad libitum. Animal experiments use and care were performed in compliance with the regulations and guidelines of Zhejiang University, and the experimental procedures were approved by the Institutional Animal Care and Research Committee of Zhejiang University (Approval #ZJU20210252 to HYF). The primers used for genotyping are listed in Supplementary Table S10.

### Superovulation and fertilization

For superovulation, female mice (28–35 days of age) were intraperitoneally injected with 5 IU of pregnant mare serum gonadotropin (PMSG; Ningbo Sansheng Pharmaceutical). After 44 h, the ovaries were punctured with needles to collect GV oocytes, or the mice were injected with 5 IU human chorionic gonadotropin (hCG, Ningbo Sansheng Pharmaceutical). After an additional 16 h, the cumulus-oocyte complex masses were surgically removed from the oviducts, and MII oocytes were collected by digestion with 0.3% hyaluronidase (Sigma-Aldrich). Oocyte morphology was observed, and images were acquired using a Nikon SMZ1500 stereoscope. To collect early embryos, 4-week-old female mice were mated with adult WT males immediately after hCG injection. Successful mating was confirmed by the presence of vaginal plugs. Embryos were harvested from the oviducts after hCG injection at the indicated time points.

### Mouse oocyte and embryo *in vitro* culture and collection

Fully grown oocytes were harvested from female mice at 28–35 days of age that were superovulated 44 h later by injection of 5 IU of PMSG in 37 °C pre-warmed M2 medium (M7167; Sigma-Aldrich) and cultured in M16 medium (M7292; Sigma-Aldrich) covered with mineral oil (M5310; Sigma-Aldrich). Mice at 28 days of age were superovulated, fertilized with adult WT males, and inspected for the presence of vaginal plugs 20 h after hCG injection. In some experiments, the obtained embryos were further cultured at 37 °C with a 5% CO_2_ atmosphere in KSOM medium (Millipore).

### Preparation of mRNA for microinjections

To prepare mRNA for microinjection, the expression vectors were linearized and *in vitro* transcribed using the SP6 message mMACHINE Kit (Invitrogen, AM1340). Poly(A) tails (∼200–250 bp) were added to the transcribed mRNAs using a mMACHINE kit (Invitrogen, AM1350). The *in vitro* transcribed mRNAs were recovered using lithium chloride precipitation and resuspended in nuclease-free water. The concentration of all injected RNAs was adjusted to 500 ng/μL. To add ac^4^C to the transcription products, acetylated cytosine and cytosine were added to the *in vitro* transcription reactions at 1:10.

### Microinjection of oocytes

Fully grown oocytes at the GV stage were incubated in an M2 medium containing 2 μM milrinone to inhibit spontaneous GV breakdown for later microinjection. All injections were administered using an Eppendorf Transferman NK2 micromanipulator. Approximately 5–10 pL of RNA at a concentration of 500 ng/µL was microinjected into each oocyte. After microinjection, oocytes were cultured in M16 medium with (arrested at the GV stage) or without (released to undergo meiotic resumption) 2 μM milrinone at 37 °C and 5% CO_2_.

### Immunofluorescence and confocal microscopy

For immunofluorescence staining, oocytes or embryos were fixed at the indicated time points in phosphate-buffered saline (PBS)-buffered 4% paraformaldehyde for 30 min at room temperature (RT) and subsequently transferred to PBS-buffered 0.3% Triton X-100 for 15 min for permeabilization. After blocking in PBS-buffered 1% bovine serum albumin, oocytes were incubated with primary antibodies diluted in blocking solution at RT for 1 h or 4 °C overnight. Oocytes were washed thrice in PBS, probed at RT with secondary antibodies for 30 min, and counterstained with 5 μg/mL of 4′,6-diamidino-2-phenylin-dole (DAPI) or propidium iodide (Molecular Probes, Life Technologies) for 10 min. Finally, the oocytes were washed and mounted on glass slides using SlowFade Gold Antifade Reagent (Life Technologies). Imaging of the oocytes or embryos after immunofluorescence was performed using a Zeiss LSM710 confocal microscope. The antibodies used in this study are listed in Supplementary Table S11. Semi-quantitative analysis of the fluorescence signals was performed using ImageJ software.

### Detection of protein synthesis in embryos

To detect nascent protein synthesis, 2-cell-stage embryos of WT and *Nat10^fl/fl^; Gdf9-Cre* were cultured in KSOM medium supplemented with 50 μM HPG from Click-iT^®^ HPG Alexa Fluor Protein Synthesis Assay kits (Thermo Fisher Scientific) for 2 h. After washing thrice with PBS, the embryos were fixed in 4% formaldehyde for 30 min at RT. After permeabilization and washing according to previously described standard protocols, Alexa Fluor 488 was conjugated to the protein using a Click-iT® cell reaction kit, and DAPI was counterstained. The fluorescence signal was subtracted from the background and measured and quantified using ImageJ software.

### Histological analysis and immunohistochemistry

Freshly isolated ovary samples extracted from WT and *Nat10^fl/fl^; Gdf9-Cre* female mice were collected and fixed overnight in PBS-buffered formalin at 4 °C, dehydrated, processed, and embedded in paraffin using standard protocols. Ovaries were serially sectioned 5 micrometers thick and stained with hematoxylin and eosin following standard protocols. For immunohistochemistry staining, sections were deparaffinized and rehydrated. The slides were then incubated in 3% H_2_O_2_ (v/v) for 10 min at RT, boiled for 15 min in 10 mM sodium citrate buffer (pH 6.0), and incubated in a blocking buffer containing 10% donkey serum for 40 min at RT. Primary antibodies were applied at suitable dilutions at 4 °C overnight, and samples were washed with PBST three times for 10 min each time and incubated with biotinylated secondary antibodies for 30 min at RT. After several washes in PBST, the sections were stained using Vectastain ABC and DAB peroxidase substrate kits (Vector Laboratories). Finally, the slides were counterstained with hematoxylin before mounting. The sections were imaged using a bright-field microscope. The antibodies used in this study are listed in Table S3.

### Western immunoblotting analysis

Protein samples of oocytes or embryos were prepared by lysing and denaturing in 2X SDS sample buffer containing β-mercaptoethanol for total protein extraction and heated for 15 min at 95°C. Proteins were separated using sodium dodecyl-sulfate polyacrylamide gel electrophoresis and electrophoretically blotted onto microporous polyvinylidene fluoride membranes (Millipore) under a constant current. Membranes containing the transferred proteins were rinsed and blocked in 0.1% Tween-20 in Tris-buffered saline (TBST) containing 5% nonfat milk (BD Biosciences) at RT for 1 h. After being probed with primary antibodies at a predetermined concentration overnight at 4°C, the membranes were washed in TBST for 15 min three times, and incubated with the corresponding horseradish peroxidase-linked secondary antibody (Jackson ImmunoResearch Laboratories) for 40 min at RT, and subsequently washed three times with TBST. Finally, the exposed bound signals were detected using an enhanced chemiluminescence western blotting substrate (Thermo Fisher Scientific, 32106) or SuperSignal West Femto maximum sensitivity substrate (Thermo Fisher). The antibodies used in this study are listed in Supplementary Table S11, and all the unprocessed gel figures were shown in Supplementary Figure S3.

### RNA isolation and real-time RT-PCR

Five oocytes or embryos were collected and lysed in 2 μL of lysis buffer (0.2% Triton X-100 and 4 IU RNase inhibitor), and cDNA synthesis was followed by retrotranscribing reverse transcription with primer transcript II reverse transcriptase (Takara) according to the manufacturer’s protocol. Quantitative RT-PCR analysis was performed using Power SYBR Green PCR Master Mix (Applied Biosystems, Life Technologies) and an Applied Biosystems ABI 7500 Real-Time PCR system using the primers listed in the supplementary material. Relative mRNA expression levels were calculated to the levels of endogenous glyceraldehyde 3-phosphate dehydrogenase mRNA (used as a housekeeping gene) using Microsoft Excel, and each RT-qPCR experimental reaction was performed in triplicate.

### LACE-seq

The LACE-seq experiments were performed according to a previously published protocol ^26^. Briefly, 50 oocytes were collected in 1.5-mL Eppendorf LoBind microcentrifuge tubes and quickly spun down to the bottom with a mini-centrifuge. Samples were lysed on ice using 50 μL of wash buffer for 10 min. Next, 1 μL of SUPERase·In RNase inhibitor (Ambion, AM2696) and 4 μL of RQ1 DNase (Promega, M6101) were applied to the lysate and incubated at 37 °C for 3 min. After snap-chilling the tube on ice for 3 min, 2 μg of ac4c antibody were added, and the tube was rotated at 4 °C for 1 h. Next, the samples were irradiated twice with UV-C light on the ice at 400 mJ, then 10 μL protein A/G beads were added to the sample and rotated for 2 h at 4 °C. After extensive washing steps, the immunoprecipitated RNA was fragmented and experienced 3′-dephosphorylation and linker ligation, followed by reverse transcription on beads. Then, first-strand cDNAs were derived from protein A/G beads and further captured by streptavidin C1 beads for pre-PCR and added 3′-cDNA linker to produce dsDNA as the template for *in vitro* transcription. *In vitro* transcription products were purified by removing the DNA template using Turbo DNase and further purified using Agencourt RNA Clean beads, according to the manufacturer’s instructions. The linearly amplified RNA was then transformed into cDNA and amplified using PCR using P7 and barcoded P5 index primers. Final PCR products of 250–500 bp were excised from a 2% agarose gel and purified using a gel purification kit (Qiagen, Catalog # 28604) according to the manufacturer’s instructions.

### LACE-seq data mapping

Adapter sequences and low-quality bases of the raw reads were removed using Trim-Galore (v.0.6.7) with the following parameters: --clip_R2 10 --clip_R1 10 -- three_prime_clip_R1 4 --three_prime_clip_R2 4 --paired --phred33 --trim-n --stringency 3 -- length 25 --fastqc. Clean reads were first aligned to mouse pre-rRNA using the Bowtie software (v.2.3.5.1) ^44^, and the remaining unmapped reads were then aligned to the mouse (mm10) reference genome using hisat2 (v.2.2.1) ^45^. For LACE-seq data mapping, two mismatches were allowed (Bowtie parameters: --sensitive-local -N 1 --local --no-mixed --un-conc-gz -p 30 -x reference genome -1 -2; Hisat2 parameters: -k 1 --no-discordant -p 30 --dta-cufflinks -x reference genome -1 -2). Then, the PCR duplicates were removed using Sambamba (v.0.7.1) ^46^. The correlation between LACE-seq replicates was calculated as follows: the LACE-seq correlation was calculated using deepTools (v.3.5.1) ‘plotCorrelation’ function with the parameters:–corMethod spearman --colorMap bwr --plotNumbers -p heatmap, then using deepTools bamcoverage with default parameters to generate bigwig files for visualization ^47^.

### LACE-seq peak and motif identification

Macs (v.2.2.7.1) software was used to identify peaks in WT and Nat10 knockout oocytes. The parameters were as follows: --keep-dup all --fe-cutoff 2 -p 0.001 --extsize 100 –nomodel^48^. Only peaks that appeared in both replicates were retained. For motif analysis, LACE-seq peaks were first extended 30 nt to 5′ upstream, and its corresponding sequence on the mm10 reference genome was obtained using bedtools (v.2.30.0) with parameters: -fi -bed -fo ^49^. Then, the Meme (v.4.11.2) software obtained their motif logo with default parameters.

### RNA-seq library preparation

For RNA-seq library construction, three replicates of zygote and 2-cell stage samples (10 embryos per sample) were collected from WT and *Nat10^fl/fl^;Gdf9-Cre* mice. Embryonic mRNA extraction and reverse transcription were performed using the Smart-seq2 protocol as previously described. Briefly, 4-week-old female mice were injected with hCG 44 hours after PMSG injection and mated with WT male mice. Embryos were collected 20 h later *in vivo*. Each sample was directly lysed in 4 μL of lysis buffer (including 0.2 μL of 1:1000 diluted external RNA controls consortium (ERCC) spike-in) and immediately reverse-transcribed using the PrimeScript^TM^ II Reverse Transcriptase (Takara, Cat# 2690A), and the cDNA library was constructed as the published Smart-seq2 method. Raw reads were sequenced using the Illumina NovaSeq 6000 platform in the 151 bp paired-end mode.

### RNA-Seq data analysis

Raw reads were trimmed with Trim-Galore v0.6.7 and mapped to the mm10 genome using STAR v2.7.10a. Uniquely mapped reads were used to quantify gene expression using FeatureCounts v2.0.2 and further normalized to the ERCC spike-in. The ERCC table was obtained as described above using the ERCC reference genome, and the percentage of ERCC reads was used for data calibration. Differential gene expression analysis was performed using the DESeq2 R package, and an adjusted P-value of < 0.05 and an absolute Log2(fold change of cKO/WT) > 1 were used as statistical significance to identify differentially expressed genes (listed in Supplementary Tables S9). Transcripts per million were calculated to estimate gene expression levels, normalized to gene length and sequencing depth. ERCC-calibrated counts and transcripts per million are listed in Supplementary Table S8.

### Statistical analysis

The experiments were randomized and performed with blinding of the experimental conditions. No statistical method was used to predetermine the sample size. Informed consent was obtained from all the subjects. Results are given as means ± SEM. Each experiment included at least three independent samples and was repeated at least three times. The results of the two experimental groups were compared using two-tailed unpaired Student’s t-tests. Statistically significant values were *P < 0.05, *P < 0.01, and *P < 0.001.

### Data availability

RNA-seq and ac^4^C LACE-seq raw data have been deposited in the NCBI Gene Expression Omnibus database (https://www.ncbi.nlm.nih.gov). The RNA-seq raw data is under accession code with GSE254288. The ac^4^C and NAT10 LACE-seq data created in this study are available from the GEO database, and the accession number is GSE253976.

## Acknowledgments

This work was supported by the National Key Research and Development Program of China (2021YFC2700100), Key Research and Development Program of Zhejiang Province (2021C03100 and 2021C03098), National Natural Science Foundation of China (31930031, 32072939), Natural Science Foundation of Zhejiang Province (LD22C060001), National Ten Thousand Talent Program, the fellowship of China National Postdoctoral Program for Innovative Talents (BX20230031), the fellowship of China Postdoctoral Science Foundation (2023M740143), and supported by Beijing Natural Science Foundation (7244435).

## Conflict of Interest

The authors declare no conflict of interest.

## Supplementary Materials

**NAT10-mediated mRNA N4-acetylation is Essential for the Translational Regulation During Oocyte Meiotic Maturation in Mice**

## Supplementary Figures

Figure S1. Comparative analyses between ac^4^C LACE-seq and acRIP-seq technologies.

Figure S2. *Nat10* deletion causes partial transcript deregulation in GV-stage oocytes.

Figure S3. Unprocessed WB gel figures.

## Supplementary Tables S1-S11 contains the following tables

Table S1. Sample correlation of LACE-seq

Table S2. ac^4^C LACE-seq FPKM-Rep 1

Table S3. ac^4^C LACE-seq FPKM-Rep 2

Table S4. ac^4^C peak annotation

Table S5. NAT10 LACE-seq FPKM Rep 1

Table S6. NAT10 LACE-seq FPKM Rep 2

Table S7. NAT10 peak annotation

Table S8. WT and *Nat10*-cko GV RNA seq TPM list

Table S9. WT and *Nat10-*cko GV RNA seq DEG list

Table S10. List of primer sequences related to experimental procedures

Table S11. Antibodies used in this study

## Supplementary Figures

**Figure S1.**
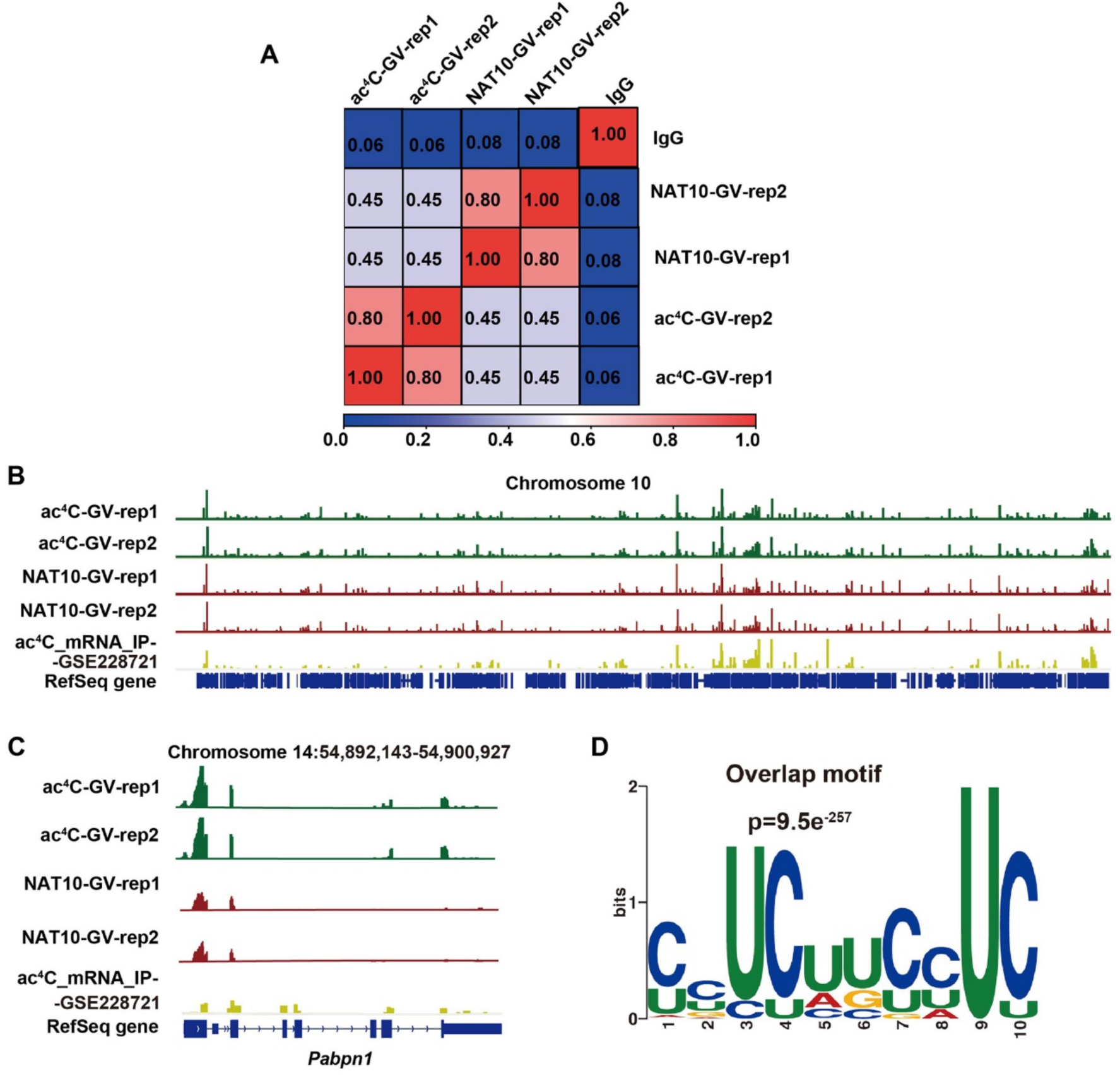
Comparative analyses between ac^4^C LACE-seq and acRIP-seq technologies. **A:** Pearson correlation coefficient between biological replicate samples. **B:** The overall distribution of peaks of chromosome 10 obtained by ac^4^C LACE-seq, NAT10 LACE-seq, and acRIP-seq. **C:** IGV results showing the ac^4^C peaks on the *Pabpn1* gene captured by ac^4^C LACE-seq, NAT10 LACE-seq and acRIP-Seq. **D:** Motif enrichment analysis of 2227 ac^4^C peaks co-targeted by ac^4^C LACE-seq and NAT10 LACE-seq.

**Figure S2.**
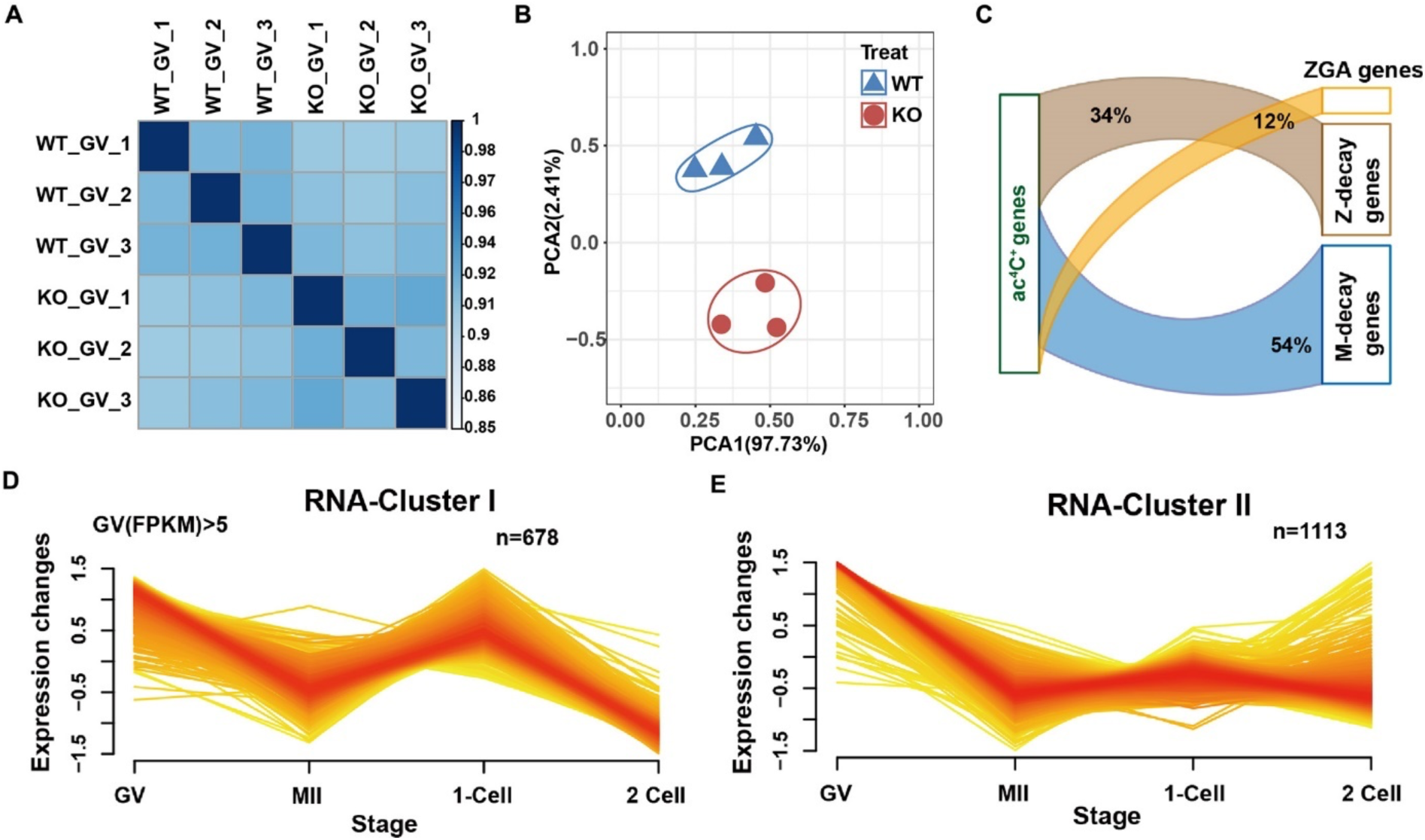
*Nat10* deletion causes partial transcript deregulation in GV stage oocytes. **A:** Heatmap showing Pearson correlation coefficients of the total transcripts between WT and *Nat10*-cKO oocyte at the GV stage. **B:** Principal component analysis (PCA) results of GV oocyte in WT and *Nat10*-cKO mice. Each symbol represents a biological sample. The proportions of variation in PCA1 and PCA2 were 97.73% and 2.41%, respectively. **C:** The ac^4^C^+^ gene was compared and analyzed with the M-decay gene, Z-decay gene, and ZGA gene in WT mice. **D-E:** The ac^4^C^+^ genes were divided into two clusters based on changes in RNA expression levels during MZT in WT mice. F: Sankey diagram showing the expression pattern of transcripts at the GV and MII stages between WT and *Nat10*-cKO mice.

## Notes

### Competing Interest Statement

The authors have declared no competing interest.

